# FOXP1 negatively regulates intrinsic excitability in D2 striatal projection neurons by promoting inwardly rectifying and leak potassium currents

**DOI:** 10.1101/2020.07.21.213173

**Authors:** Nitin Khandelwal, Sheridan Cavalier, Volodymyr Rybalchenko, Ashwinikumar Kulkarni, Ashley G. Anderson, Genevieve Konopka, Jay R. Gibson

**Affiliations:** Department of Neuroscience, University of Texas Southwestern Medical Center, Dallas, TX, 75390, USA

## Abstract

Heterozygous loss-of-function mutations in the transcription factor *FOXP1* are strongly associated with autism. Dopamine receptor 2 expressing (D2) striatal projection neurons (SPNs) in heterozygous *Foxp1* (*Foxp1*^+/−^) mice have higher intrinsic excitability. To understand the mechanisms underlying this alteration, we examined SPNs with cell-type specific homozygous *Foxp1* deletion to study cell-autonomous regulation by *Foxp1*. As in *Foxp1*^+/−^ mice, D2 SPNs had increased intrinsic excitability with homozygous *Foxp1* deletion. This effect involved postnatal mechanisms. The hyperexcitability was mainly due to down-regulation of two classes of potassium currents: inwardly rectifying (K_IR_) and leak (K_Leak_). Single-cell RNA sequencing data from D2 SPNs with *Foxp1* deletion indicated the down-regulation of transcripts of candidate ion channels that may underlie these currents: *Kcnj2* and *Kcnj4* for K_IR_ and *Kcnk2* for K_Leak_. This *Foxp1*-dependent regulation was neuron-type specific since these same currents and transcripts were either unchanged, or very little changed, in D1 SPNs with cell-specific *Foxp1* deletion. Our data are consistent with a model where FOXP1 negatively regulates the excitability of D2 SPNs through K_IR_ and K_Leak_ by transcriptionally activating their corresponding transcripts. This, in turn, provides a novel example of how a transcription factor may regulate multiple genes to impact neuronal electrophysiological function that depends on the integration of multiple current types – and do this in a cell-specific fashion. Our findings provide initial clues to altered neuronal function and possible therapeutic strategies not only for FOXP1-associated autism but also for other autism forms associated with transcription factor dysfunction.

## Introduction

Forkhead box P1 (FOXP1) is a transcription factor that is highly enriched in the developing embryonic and mature neocortex, hippocampus, and striatum (1). Heterozygous mutations and deletions in *FOXP1* resulting in loss-of-function are causative for Intellectual Disabilities (ID) and Autism Spectrum Disorders (ASD) (2–4) and are among the most significant recurrent *de novo* mutations associated with ASD (5–7). Characterization of whole brain *Foxp1* conditional knockout mice (cKO) has shown that these mice exhibit ASD-relevant repetitive behavior and reduced social interaction (8). Moreover, heterozygous *Foxp1* (*Foxp1*^+/−^) mice, which are a relevant genetic model for *FOXP1*-associated ASD, also show altered vocalizations (9).

ASD patients have altered striatal function and connectivity (10–13), and *Foxp1* is among the few genes specifically enriched in the mouse striatum (14). Deletion of *Foxp1* in the whole brain leads to a dramatic reduction in striatal size and is required for striatal projection neuron (SPN) differentiation *in vitro*, indicating its critical role in striatal development (8, 9, 15, 16). Beginning at embryonic stages and continuing into adulthood, *Foxp1* is expressed in dopamine receptor subtype 1 (D1) and dopamine receptor subtype 2 (D2) expressing SPNs (1, 9, 16, 17). However, very little is known about the role of *Foxp1* in the maturation of these striatal neurons.

Some mechanistic details, such as the role of SUMOylation, in FOXP1-dependent regulation of brain development have been reported in the past (18). However, none of the studies have made a compelling link between downstream targets of FOXP1 – either direct or indirect – with neuronal development. Moreover, previous studies have mostly employed embryonic *Foxp1* deletion and examined early developmental processes (19–22), but as *Foxp1* is expressed in adult brain as well, its involvement in postnatal development is also possible. These gaps in understanding apply to other members of the FOXP family, like FOXP2. Therefore, further examination of FOXP1 function is needed, not only for FOXP1 alone, but also for the FOXP family as a whole.

FOXP1 has been reported to be involved in the development of synaptic transmission and intrinsic excitability – the latter being the propensity to fire action potentials (8, 9, 23). Brain-wide conditional *Foxp1* deletion decreases intrinsic excitability of hippocampal pyramidal neurons (8). In *Foxp1*^+/−^ mice, intrinsic excitability is increased in D2 SPNs, but not in D1 SPNs (9). These studies indicate that FOXP1 may have differing roles depending on neuron type, but this issue is confounded by the different mouse models used and has not been addressed so far. Furthermore, mechanistic details of these excitability alterations remain unknown.

Both D1 and D2 SPNs express several potassium (K^+^) channels, including two classes of subthreshold K^+^ channels – leak (K_Leak_) and inwardly rectifying (K_IR_) (24–29). K_Leak_ channels are the two-pore domain channels that are not gated by membrane potential (30). Evidence supports the existence of K_Leak_ channels, particularly KCNK2 (TREK-1), in SPNs (28, 29, 31, 32). K_IR_ currents in SPNs are mediated by KCNJ2 (Kir2.1) and KCNJ4 (Kir2.3) channels (24, 25, 27, 33) and are a major source of conductance at hyperpolarized potentials. While some signaling pathways in SPNs have been reported to play a role in the maturation of intrinsic excitability through the regulation of K_IR_ currents (27, 34), the transcriptional regulation of this maturation is not known.

Transcriptional regulation of neuronal function during development, and its impairment in ASD, is not well understood. The difficulty is due to the large number of genes impacted by transcriptionally-related proteins and to differing roles in different cell-types. Among the top 50 most commonly studied autism-associated genes (35), 14 are involved in transcriptional regulation. Among these, which includes *Foxp1*, a connection between ion channel transcripts and function has only been made for *Mef2c* and *Mecp2* (36, 37). And for both of these, direct demonstration on how this link affects neuronal intrinsic excitability has not been reported. And for both, a description of a more multi-faceted regulation involving multiple downstream transcripts and corresponding channel function and how this contributes to neuronal excitability in different neuron-types was lacking.

Using a cell-autonomous deletion strategy in either D1 or D2 SPNs, we examine the electrophysiological mechanisms by which FOXP1 regulates the maturation of intrinsic excitability. In D2 SPNs, we find that *Foxp1*-dependent regulation involves multiple currents and transcripts. However this regulation is either absent or diminished in D1 SPNs. Specifically, we find that FOXP1 negatively regulates hyperexcitability by promoting the function of K_IR_ and K_Leak_ currents in D2 SPNs during development. Using striatal singlecell RNA sequencing data, we also identify specific candidate channels that underlie these currents: KCNJ2 and KCNJ4 for K_IR_ and KCNK2 for K_Leak_.

## Materials and Methods

### Mice

All mice used were of C57BL/6J background strain. *Drd2* conditional *Foxp1* knockout (D2 *Foxp1^cKO^*) mice were: *Drd2*-Cre^tg/-^:*Foxp1*^flox/flox^:*Drd2*-eGFP^tg/-^, while *Drdlα Foxp1^cKO^* (D1 *Foxp1^cKO^*) were: *Drdlα*-Cre^tg/-^:*Foxp1*^flox/flox^:*Drdlα*-tdTomato^tg/-^. Heterozygous version of D2 *Foxp1^cKO^* is referred by D2 *Foxp1^cHet^*.

### Brain slices and Recordings

Unless stated otherwise, P14-23 mice were used for the electrophysiological recordings. Only D2 SPNs were recorded in D2 *Foxp1^cKO^* mice, where WT controls and *Foxp1*-deleted neurons were named *Foxp1^CTL^* and *Foxp1^cKO^* D2 SPNs, respectively. Similar nomenclature applies to D1 SPNs. For the slices where *Foxp1* was deleted postnatally using virus, uninfected SPNs were named *Foxp1^CTL^* SPNs while virus-infected neurons were named *Foxp1^vcKO^* SPNs.

### Current steps

In current clamp, incremental current steps (500 ms duration) were applied at resting potential to measure number of action potentials elicited at each step to make F-I curve.

### Single −10 voltage step to measure input resistance, capacitance, and conductance

In voltage clamp, a single −10 mV voltage step (500 ms duration) was applied to measure input resistance, capacitance, normalized cell conductance, and the voltage-dependence of holding current at −85, −65, and − 55 mV holding potentials.

### Multi-step protocol using IV-plots

In voltage clamp, a multi-step protocol (500 ms duration) was applied while maintaining −60 mV holding potential (ranging from −30 to −120 mV voltages in −10 mV steps). The current measured was the average of a 200 ms window at the end of the step (see Fig. 2E).

### Morphology and analysis

Dendritic morphology was measured by filling neurons with a fluorescent dye (AF488, 100 μM, Life Technologies, Inc.) and biocytin (2 mg/ml, Life Technologies, Inc.) during the whole-cell recording. Sholl analysis is performed using concentric radii step intervals of 20 μm and dendritic crossings at each radius are counted.

### Statistics

For all data, sample number is the number of neurons, and data are stated in this order: WT control followed by *Foxp1* deletion groups. All data are plotted as mean ± standard error. Unless stated otherwise, a repeated measures (RM) 2-way ANOVA is performed with Geisser-Greenhouse correction followed by the Holm-Sidak test for the correction for multiple comparisons. When ANOVA analysis is not employed, we use a Mann-Whitney (MW) t-test.

### Single-cell RNA sequencing

We used single-cell RNA sequencing data from our previous study and analyzed D1 and D2 SPNs (15).

## Results

### Increased intrinsic excitability in D2 SPNs with cell autonomous Foxp1 deletion

We first examined how intrinsic excitability was altered in D2 SPNs with homozygous *Foxp1* deletion. We performed selective *Foxp1* deletion in D2-SPNs using D2 *Foxp1^cKO^* mice to isolate FOXP1-regulated cell-autonomous mechanisms. We targeted D2 SPNs by their GFP expression and obtained “firing versus current injection” curves, or “F-I” curves (Fig. 1A,B). *Foxp1^cKO^* D2 SPNs exhibited a ~30-500% increase in the number of evoked action potentials when compared to the *Foxp1^CTL^* D2 SPNs, indicating that FOXP1 negatively regulates intrinsic excitability of D2 SPNs.

**Figure 1.**
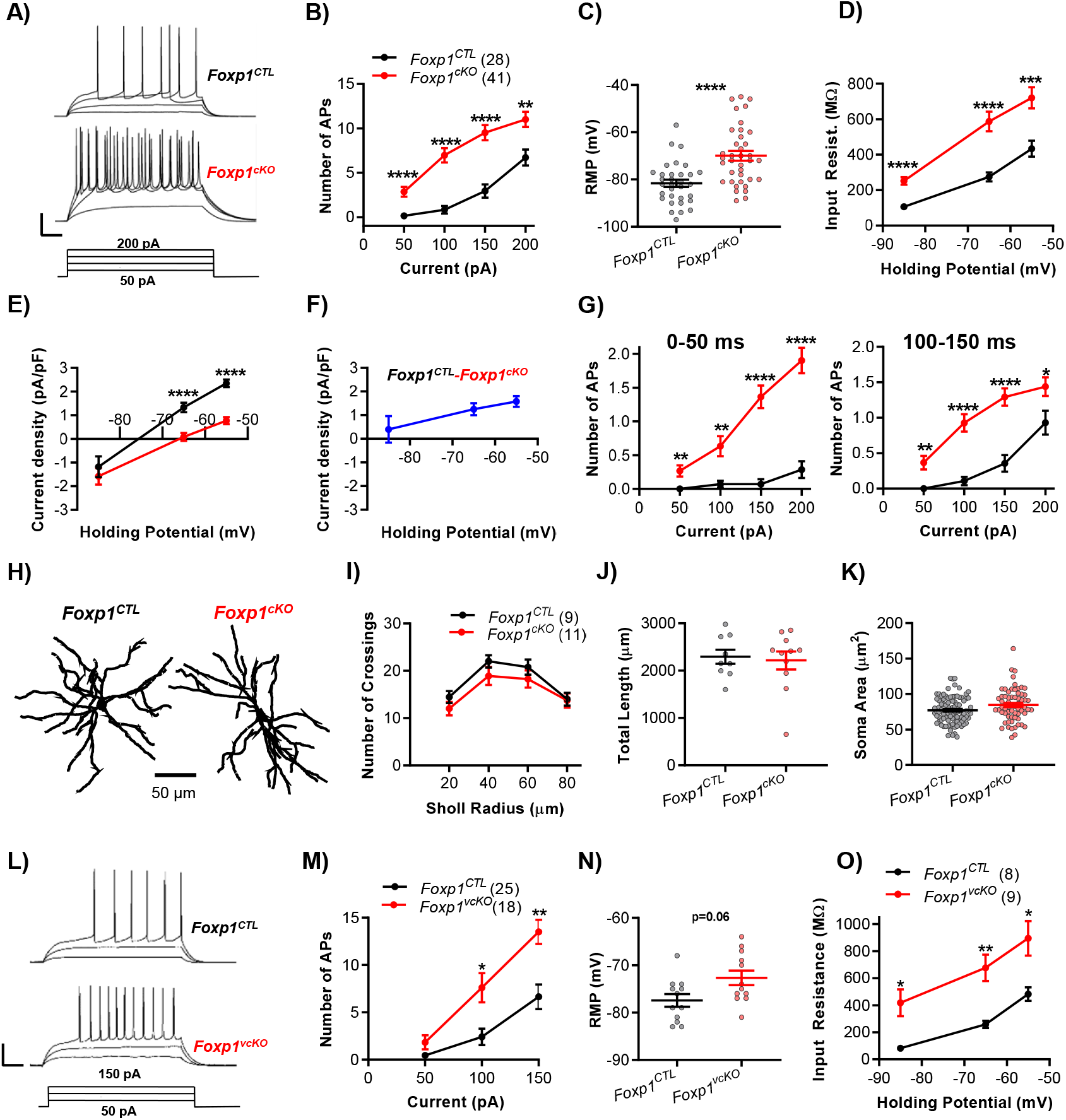
Cell-autonomous *Foxp1* deletion leads to increased intrinsic excitability in D2 SPNs but does not affect their morphology. A) Example traces of action potentials evoked by depolarizing current steps. Scale bars: 60 ms, 30 mV. B) Action potential (AP) number versus current step amplitude curves (F-I curves) indicate increased number of spikes in *Foxp1^cKO^* SPNs. n’s in B also apply to D-G. C) Resting membrane potential (RMP) was significantly higher in *Foxp1^cKO^* SPNs, n=32,37. D) Input resistance at - 85 mV, −65 mV and −55 mV holding potentials was elevated in *Foxp1^cKO^* SPNs. E) Current density is plotted as a function of holding potential. F) The difference plot based on the plots in E illustrates the down-regulated current in *Foxp1^cKO^* SPNs. G) F-I curves in the first 50 ms (*left*) immediately after the onset and during the 100-150 ms (*right*) window of the current step. H) Example of dendritic reconstructions of D2 SPNs. I) Sholl analysis revealed no change in the number of dendritic branches with *Foxp1* deletion. J,K) No difference was observed in the total dendritic length (J, n=9,11) and soma area (K, n=94,63). L) Example traces of action potentials evoked by depolarizing current steps in *Foxp1^CTL^* and *Foxp1^vcKO^* D2 SPNs (deletion induced with Cre-expressing virus). Scale bars: 60 ms, 30 mV. M) F-I curves indicate increased intrinsic excitability of AAV-Cre infected D2 SPNs. N) A trending increase in resting membrane potential was observed in *Foxp1^vcKO^* D2 SPNs compared to uninfected controls (*Foxp1^CTL^*) (N=12,12). O) Input resistance was increased in *Foxp1^vcKO^* D2 SPNs at all the three membrane potentials. Statistical analysis: For B, D, E, G, I, M and O, RM 2W-ANOVA. As in all figures using ANOVA, asterisks indicate post-hoc comparisons. For C, J, K and N, M-W t-test. *p<0.05, **p<0.01, ***p<0.001, ****p<0.0001.

Subthreshold membrane properties of the D2 SPNs were also altered upon *Foxp1* deletion as observed by higher resting membrane potential (Fig. 1C), higher input resistance (Fig. 1D), and altered holding current density (Fig. 1E). When the average holding current density in *Foxp1^cKO^* D2 SPNs was subtracted from that of *Foxp1^CTL^* D2 SPNs, a difference current density was obtained with a reversal potential near −85 mV (Fig. 1F). This is very close to the estimated reversal potential for K^+^ in our experiments (−88 mV). These changes indicate that a down-regulation of subthreshold K^+^ currents with *Foxp1* deletion causes an increase in input resistance and a depolarized resting potential, thereby contributing to their hyperexcitability.

To examine how these subthreshold alterations contributed to increased intrinsic excitability, we re-analyzed the F-I data in figure 1B and plotted the number of spikes at different epochs during the current step (Fig. 1G). Action potential firing in the first 50 ms epoch is expected to depend most on the subthreshold resting state. For the subsequent epochs, depolarization-activated currents are engaged and the impact of the previous subthreshold properties is diminished. The greatest increase in excitability is observed in the first 50 ms epoch compared to all other subsequent epochs suggesting that subthreshold membrane properties are involved in the hyperexcitability of *Foxp1^cKO^* D2 SPNs.

To better isolate the role of input resistance and spike voltage-threshold changes underlying increased intrinsic excitability, we analyzed the current step data only for the neurons having resting potential in the range between −65 mV to −85 mV. The F-I curves showed similar hyperexcitability for *Foxp1^cKO^* D2 SPNs (Data not shown). In this analysis, the average resting potential and spike voltage-threshold were not different in *Foxp1^cKO^* D2 SPNs (Table 1). However, spike current-threshold and input resistance were both approximately doubled. Therefore, increased input resistance appears to be a key mediator for hyperexcitability.

To determine if D2 SPN hyperexcitability in *Foxp1*^+/−^ mice is due to cell-autonomous deletion (9), we performed the same experiments in D2 SPNs that were selectively heterozygous for *Foxp1* (*Foxp1*^cHet^). We found no change in excitability or subthreshold properties (Supplementary Fig. S1A-D) indicating that hyperexcitability in D2 SPNs of the *Foxp1*^+/−^ mouse may not be due to a purely cell-autonomous process.

### No significant morphological or anatomical changes with Foxp1 deletion in D2 SPNs

It is possible that *Foxp1^cKO^* D2 SPNs are smaller in size and the resulting reduction in membrane surface area could contribute to increased input resistance and increased excitability. To test this idea, we filled D2 SPNs with biocytin, and measured branching complexity (Sholl Analysis) and total dendritic length. We could not find any effect of *Foxp1* deletion on these properties (Fig. 1H-J), which suggests that cell morphology is not affected. We also measured soma size in live, acutely prepared slices and found no difference (Fig. 1K). In summary, there was no anatomical evidence for change in membrane surface area. Another parameter that crudely estimates the membrane surface area is the cell capacitance. When pooled over all experiments conducted under control conditions in our study, there was no detectable change in the cell capacitance after *Foxp1* deletion (Supplementary Fig. S2, *Control*). However, under strong blockade of multiple channels, which may more accurately report membrane surface area (or “cell size”) due to better space clamp, *Foxp1^cKO^* D2 SPNs had a 19% reduction in capacitance (Supplementary Fig. S2, *Cs^+^,Ba^2+^*). In summary, while this change is difficult to interpret, there may be some contribution of decreased membrane surface area to their hyperexcitability.

### Postnatal Foxp1 deletion in D2 SPNs also results in hyperexcitability

It is unclear whether hyperexcitability of D2 SPNs is due to deletion of *Foxp1* during embryonic or postnatal development. When *Foxp1* is deleted during embryonic development, indirect effects involving cell fate determination and migration may affect excitability. This could indeed be a factor since D2 SPN number is reduced in D2 *Foxp1^cKO^* mice (see Suppl. Methods). However, if hyperexcitability can be induced with *Foxp1* deletion at a postnatal stage, then the underlying mechanism is likely independent of these early developmental processes. To test this, we deleted *Foxp1* postnatally by infecting SPNs with AAV-Cre-GFP in P0-1 *Foxp1^flox/flox^* mice, resulting in *Foxp1* deletion by P10 (see Supp. Methods). Uninfected D2 SPNs in the same mice served as controls (*Foxp1^CTL^*).

The F-I curves showed an increased intrinsic excitability of D2 SPNs with *Foxp1* deletion (Fig. 1L,M). There was a statistical trend for a depolarized resting potential (p=0.06, Fig. 1N) and input resistance was higher (Fig. 1O) in *Foxp1^vcKO^* D2 SPNs, which was consistent with the down-regulation of a subthreshold K^+^ current. Dendritic complexity, total dendritic length, soma size, and capacitance were unaltered (Supplementary Fig. S3A-E). When AAV-Cre-GFP was injected into WT mice, none of the electrophysiological changes were observed (Supplementary Fig. S4A-C). These results indicate a postnatal role for *Foxp1* in regulating excitability.

With postnatal deletion, there was no decrease observed in D2 SPN number as with embryonic deletion in D2 *Foxp1^cKO^* mice (see Suppl. Methods) while the mechanism inducing hyperexcitability is similar for both deletion strategies. Since embryonic deletion was more efficient, consistent, and less intrusive, we used D2 *Foxp1^cKO^* mice for the remainder of this study.

### Down-regulation of Cs^+^-sensitive currents indicates that K_IR_ is decreased in D2 SPNs with Foxp1 deletion

To determine the identity of the subthreshold K^+^ currents that are downregulated with *Foxp1* deletion, we examined the effect of specific K^+^ channel blockers on D2 SPNs. K_IR_ current (K_IR_) is a prime candidate since K_IR_ channels are highly expressed in SPNs and are open only at hyperpolarized, subthreshold membrane potentials (24, 25, 27, 33). To measure the effect of *Foxp1* deletion on K_IR_, we used Cs^+^ to isolate K_IR_ at subthreshold potentials (25, 38–40) (Supplementary Table S1,S2).

In nominal ACSF, we observed the similar approximate doubling of the input resistance in *Foxp1^cKO^* D2 SPNs as observed in figure 1D (Fig. 2A). We also calculated normalized conductance as a measure of membrane ion permeability that is more independent of membrane surface area and found this to be roughly halved (Fig. 2C). With application of CsCl (1 mM), the effects relevant to K_IR_ were best observed at −85 mV membrane potential (see Supp. Methods, *Membrane potentials for measuring K_IR_ and K_Leak_*). For both *Foxp1^CTL^* and *Foxp1^cKO^* D2 SPNs, there was an increase in the input resistance and decrease in the normalized conductance at −85 mV (Fig. 2A,C). Consistent with the down-regulation of K_IR_ with *Foxp1* deletion, these Cs^+^-induced absolute changes at −85mV were less in *Foxp1^cKO^* D2 SPNs. However, the proportional change in both of these properties was not different between *Foxp1^CTL^* and *Foxp1^cKO^* D2 SPNs (Fig. 2B,D) indicating that Cs^+^-sensitive currents made equal proportional contributions to these properties in both groups. These results indicate that while *Foxp1* deletion did indeed reduce K_IR_, it did not alter the proportion of K_IR_ relative to other currents.

**Figure 2.**
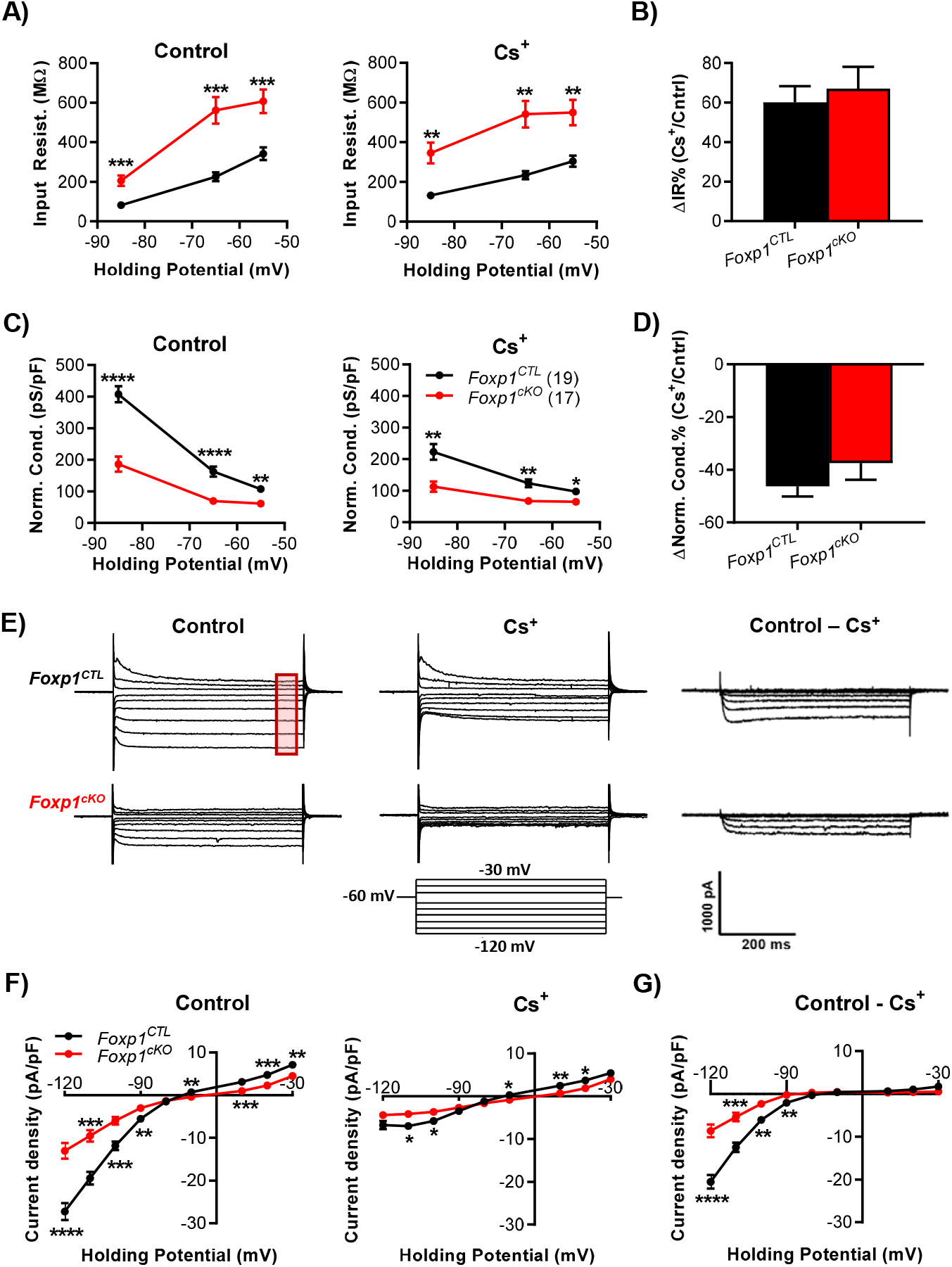
Cs^+^-sensitive inwardly rectifying K^+^ current (K_IR_) is down-regulated in D2 SPNs with embryonic *Foxp1* deletion. A) Input resistance was increased in *Foxp1^cKO^* D2 SPNs both before (*left*) and after Cs^+^-application (*right*). B) Cs^+^ exerted a similar proportional effect on the input resistance of *Foxp1^CTL^* and *Foxp1^cKO^* SPNs at −85 mV membrane potential. C) *Foxp1^cKO^* D2 SPNs had reduced normalized conductance both before (*left*) and after Cs^+^-application (*right*). n’s in C apply to all panels. D) The proportional effect of Cs^+^ was similar on the conductance of both genotypes at −85 mV membrane potential. E) Example current traces collected in the multi-step voltage protocol in control ACSF (*left*) and in Cs^+^ (*middle*) which were subtracted to obtain the Cs^+^-sensitive K_IR_ current (Difference currents, *right*). The red box indicates when current density was averaged to prepare IV-plots. F) From the multi-step protocol, average current density versus voltage plots (IV-plots) were obtained for control (*left*) and Cs^+^ application (*right*). G) Difference current IV-plots indicate reduced Cs^+^-sensitive K_IR_ current in *Foxp1^cKO^* D2 SPNs. Statistical analysis in all panels: RM 2W-ANOVA. *p<0.05, **p<0.01, ***p<0.001, ****p<0.0001.

We used a multi-step voltage protocol to measure K_IR_ current density (Fig. 2E). Current-voltage plots (IV-plots) derived from these measurements indicated that evoked current densities were consistently less in *Foxp1^cKO^* D2 SPNs compared to *Foxp1^CTL^* D2 SPNs and application of Cs^+^ affected these currents (Fig. 2E,F). For each neuron, the amount of Cs^+^-sensitive K_IR_ was approximated by subtraction of currents (Fig. 2E,*right*). Average of these difference currents indicated a significant reduction in K_IR_ with *Foxp1* deletion as observed in the steps to −120 to −90 mV (Fig. 2G). With conditional heterozygous deletion, we observed no change in K_IR_ (Supplementary Fig. S5A-C), which is consistent with the unaltered intrinsic excitability under these conditions (see Supplementary Fig. S1).

### Down-regulation of Ba^2+^-sensitive/ Cs^+^-insensitive currents indicates that K_Leak_ is decreased in D2 SPNs with Foxp1 deletion

Both K_IR_ and K_Leak_ are thought to be prominent subthreshold currents in SPNs. Because differences in subthreshold currents and conductances remained between *Foxp1^CTL^* and *Foxp1^cKO^* D2 SPNs after blocking K_IR_ with Cs^+^, we next determined if the remaining difference was due to a putative K_Leak_ (using the term “putative” to acknowledge the lack of effective, specific K_Leak_ blockers in neurons). We examined K_Leak_ through application of BaCl2 (4 mM) since barium blocks most K_Leak_ channel subtypes – including KCNK2 (Supplementary Table S1,S2). CsCl (1 mM) was always present to examine membrane properties independent of K_IR_. This experiment directly measures FOXP1-regulation of Ba^2+^-sensitive/Cs^+^-insensitive currents, and many K_Leak_ subtypes fit this description - including KCNK2. Because Cs^+^ does not completely block K_IR_ (39, 40), currents collected in control conditions at and below −75 mV membrane potential may have included some residual K_IR_, and therefore, currents at and above −65 mV membrane potential were analyzed for putative K_Leak_ (see Supp. Methods, *Membrane potentials for measuring K_IR_ and K_Leak_*).

Application of Ba^2+^ led to a decrease in conductance (Fig. 3A) and an increase in input resistance (Supplementary Fig. S6A,B) across all membrane potentials in both *Foxp1^CTL^* and *Foxp1^cKO^* D2 SPNs. *Absolute* changes in these properties were smaller in *Foxp1^cKO^* D2 SPNs which was consistent with reduced putative K_Leak_. If FOXP1 preferentially regulates K_Leak_, we would predict a decreased *proportional* effect on conductance and input resistance by Ba^2+^ application in *Foxp1^cKO^* D2 SPNs. Indeed, this was the case (Fig.3B, Supplementary Fig. S6C). From this, we conclude that *Foxp1* deletion preferentially down-regulated putative K_Leak_ compared to other subthreshold currents.

**Figure 3.**
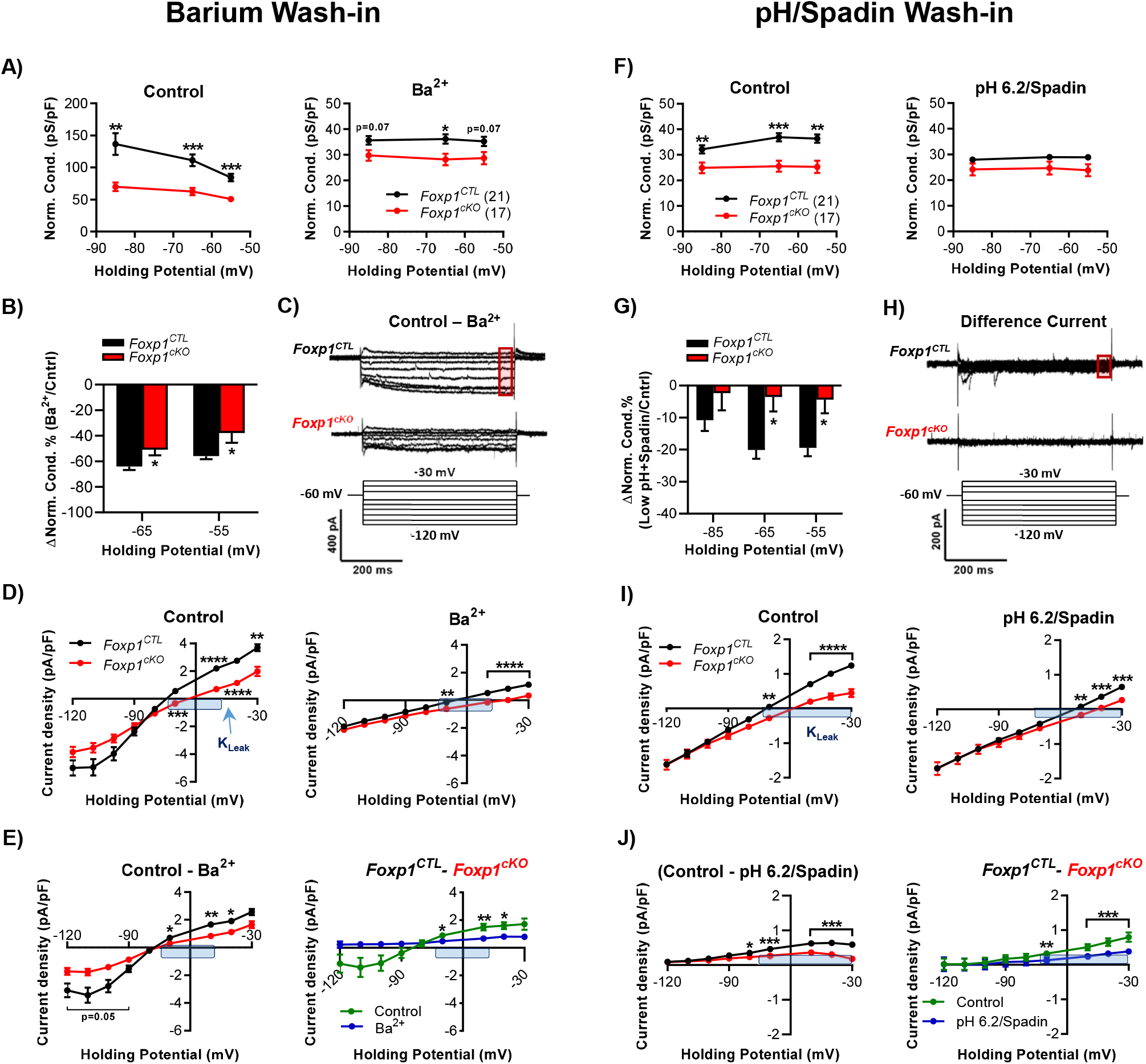
Cs^+^-insensitive K_Leak_ is preferentially down-regulated by *Foxp1* deletion. *Barium wash-in (left):* CsCl (1 mM) was present throughout all experiments. A) Conductance was decreased in both *Foxp1^CTL^* and *Foxp1^cKO^* D2 SPNs after Ba^2+^ application. n’s in A also apply to B, D and E. B) Ba^2+^ induced a lower proportional change in the conductance of *Foxp1^cKO^* D2 SPNs. C) Example current traces of the Ba^2+^-sensitive/Cs^+^-insensitive difference currents during the multi-step protocol. The red box indicates when current density was averaged to prepare IV-plots. D) IV-plots for the currents in control ACSF (*left*) and after Ba^2+^ application (*right*). E) Reagent difference IV plots (*left*) display Ba^2+^-sensitive/Cs^+^-insensitive current while genotypic difference plots (*right*) exhibit the disparity between both the genotypes for control ACSF (derived from D, *left*) and in presence of Ba^2+^ (derived from D, *right*). Blue boxes mark the region of interest for putative K_Leak_ measurements in D-E. *pH-Spadin wash-in (right):* CsCl (1 mM) and BaCl2 (4 mM) were present throughout all experiments. F) Acidification (pH8.0→6.2) and Spadin (500 nM) application noticeably reduced the conductance in *Foxp1^CTL^*, but not in *Foxp1^cKO^*, D2 SPNs. n’s in F apply also to G, I and J. G) The proportional change in conductance was lower in *Foxp1^cKO^* D2 SPNs. H) Example current traces of the difference current collected in the multi-step protocol. I) IV-plots obtained in control ACSF (*left*) showed significant differences at potentials relevant to K_Leak_ (−70 mV to −30 mV, highlighted with blue box), which substantially reduced in pH 6.2/Spadin (*right*). J) The reagent difference IV-plot (*left*) indicates that *Foxp1^cKO^* D2 SPNs have significantly reduced average difference current compared to *Foxp1^CTL^* D2 SPNs, however, this difference current is not purely K_Leak_. Genotypic difference plot (*right*) shows down-regulation of putative K_Leak_ currents in *Foxp1^cKO^* D2 SPNs as pH 6.2/Spadin-application reduced the K_Leak_ disparity between *Foxp1^CTL^* and *Foxp1^cKO^* D2 SPNs. Statistical analysis: For all panels except E (*right*) and J (*right*), RM 2W-ANOVA. (For E (*right*) and J (*right*), see Suppl. Methods, *Multi-step protocol using IV-plots*). *p<0.05, **p<0.01, ***p<0.001, ****p<0.0001.

Data from IV-plots (Fig. 3C, Supplementary Fig. S7) further supported K_Leak_ being reduced with *Foxp1* deletion. As expected for incomplete K_IR_ blockade, plots obtained in control ACSF at membrane potentials below −70 mV displayed inflections indicating that K_Leak_ was not isolated at those potentials (Fig. 3D,*left*). Application of barium strongly reduced current densities at all membrane potentials by blocking most of the K_Leak_ and residual K_IR_ and induced a linearization of the IV-plots (Fig. 3D,*right*). As depicted in difference IV-plots (Fig. 3E,*left*), the reduction in current density in *Foxp1^cKO^* D2 SPNs was less compared to *Foxp1^CTL^* D2 SPNs at depolarized membrane potentials, particularly at −70 and −50 mV, confirming that putative K_Leak_ was down-regulated with *Foxp1* deletion. The same data replotted into genotypic difference IV-plots (Fig. 3E,*right*) not only confirmed our finding, but also validated that the down-regulated current was a K^+^ current as it either crossed at or was not detectably different from the x-axis at the K^+^ reversal potential (−88 mV).

We also examined the down-regulation of Ba^2+^- and Cs^+^-sensitive currents combined. In the earlier Cs^+^-application experiment, the disparity in conductance between *Foxp1^CTL^* and *Foxp1^cKO^* D2 SPNs in control conditions was 215 pS/pF (Fig. 2C,*left*, at −85 mV). When we block K_IR_ and K_Leak_ with both Ba^2+^ and Cs^+^, this disparity was reduced to 6 pS/pF (Fig. 3A,*right*, at −85 mV). In terms of percentage, the conductance disparity decreased from 53% to 17% of control. Therefore, K_Leak_ and K_IR_ account for a large portion of the conductance that is down-regulated with *Foxp1* deletion.

### Targeting KCNK2 channels further implicates K_Leak_ down-regulation with Foxp1 deletion

Pharmacological blockade in the experiments above did not provide a complete accounting for currents down-regulated with *Foxp1* deletion since differences remained between *Foxp1^CTL^* and *Foxp1^cKO^* SPNs (Fig. 3A,*right, 3D,right*). One possible reason could be an incomplete block of K_Leak_ by Ba^2+^. KCNK2 is the most expressed K_Leak_ channel in the striatum (28, 29) which is only 90% blocked by Ba^2+^ (41).

We examined whether additional blockade of KCNK2 channels could further reduce the disparity in currents between *Foxp1^CTL^* and *Foxp1^cKO^* D2 SPNs which would provide additional evidence that K_Leak_ is regulated by FOXP1. We first performed recordings in control ACSF and then in the presence of acidification (pH6.2) and Spadin (500 nM, referred to by pH6.2/Spadin-application) to specifically block KCNK2 channels (42–45). Because we wanted to determine the effect of blocking KCNK2 channels relative to the Cs^+^ and Ba^2+^ blockade examined above, extracellular Cs^+^ and Ba^2+^ were always present.

As observed in similar conditions above (see Fig. 3A,*right*), normalized conductance was significantly lower in *Foxp1^cKO^* D2 SPNs than *Foxp1^CTL^* SPNs in control ACSF (Fig. 3F,*left*). With pH6.2/Spadin-application, this disparity was no longer observed (Fig. 3F,right). Furthermore, the percent reduction in the normalized conductance was higher in *Foxp1^CTL^* D2 SPNs as compared to *Foxp1^cKO^* D2 SPNs indicating a greater contribution of pH6.2/Spadin-sensitive currents in control SPNs (Fig. 3G). These results indicated that KCNK2 was preferentially decreased with *Foxp1* deletion relative to other subthreshold currents.

In IV-plots, we observed effects of pH6.2/Spadin-application (Fig. 3H,I, Supplementary Fig. S8), and the difference IV-plot (Fig. 3J,*left*) indicated less pH6.2/Spadin-sensitive currents in *Foxp1^cKO^* D2 SPNs compared to *Foxp1^CTL^* D2 SPNs. These difference plots also indicated that pH6.2/Spadin-sensitive currents were not purely KCNK2-mediated since they did not cross the x-axis at the expected reversal potential for K^+^ and displayed a downward deflection at depolarized potentials. We attribute these effects to a nonspecific effect of acidification on membrane currents. However, when we replot the data into genotypic difference IV-plots (Fig. 3J,*right*), the non-specific effects are subtracted away and the difference currents at −70 mV and above illustrate down-regulation of KCNK2 in *Foxp1^cKO^* D2 SPNs (see Supp. Methods, *Membrane potentials*….).

### D1 SPNs become hyperexcitable with Foxp1 deletion, but through different mechanisms compared to D2 SPNs

In *Foxp1*^+/−^ mice, D2 SPNs were hyperexcitable, but not D1 SPNs (9). Therefore, either *Foxp1* is not involved in maturation of intrinsic excitability or a single copy of the *Foxp1* allele is sufficient to maintain normal excitability in D1 SPNs. To investigate this further, we examined the cell-autonomous effects of *Foxp1* deletion in D1 SPNs using D1 *Foxp1^cKO^* mice. In F-I curves, we observed increased intrinsic excitability in *Foxp1^cKO^* D1 SPNs compared with *Foxp1^CTL^* D1 SPNs (Fig. 4A,B), indicating a cell-autonomous role for FOXP1 in the proper maturation of intrinsic excitability. However, subthreshold current changes in *Foxp1^cKO^* D1 SPNs were much less than that observed in *Foxp1^cKO^* D2 SPNs. The changes in resting potential (Fig. 4C) and input resistance (Fig. 4D) were either small or not significantly different. IV-plots were unchanged – including the difference plot depicting isolated K_IR_ (Fig. 4E,F). Postnatal *Foxp1* deletion, using AAV-Cre-GFP infection in D1 SPNs, also did not induce any changes in excitability or subthreshold membrane properties (Supplementary Fig. S9A-C).

**Figure 4.**
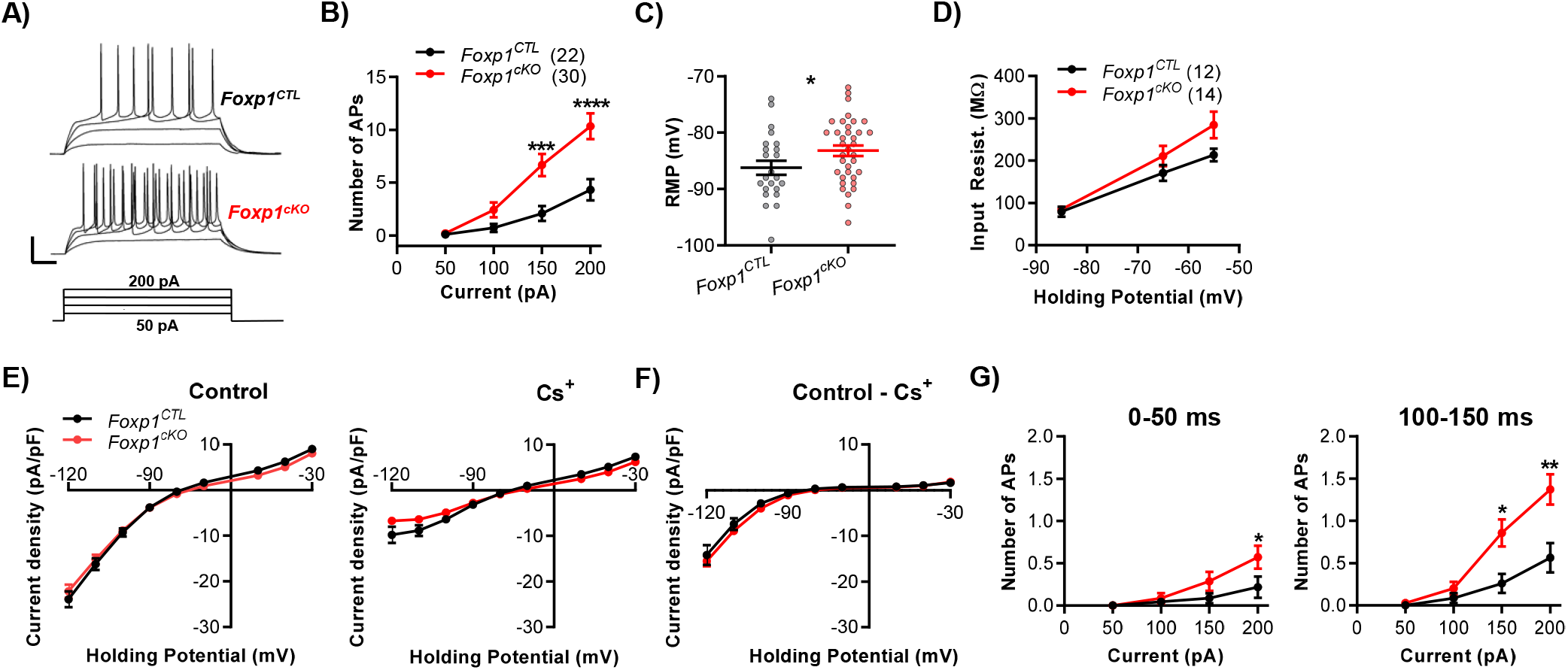
Cell-autonomous, embryonic deletion of *Foxp1* increases the intrinsic excitability in D1 SPNs, but subthreshold properties are unchanged. A) Example traces of action potentials evoked by depolarizing current steps in *Foxp1^CTL^* and *Foxp1^cKO^* D1 SPNs. Scale bars: 60 ms, 30 mV. B) F-I curves indicate increased intrinsic excitability in *Foxp1* deleted D1 SPNs. n’s in B also apply to G. C) Resting membrane potential was slightly higher in the D1 SPNs with *Foxp1* deletion. D) No change in input resistance was resolved with *Foxp1* deletion. n’s in D also apply to E and F. E,F) No difference was observed between genotypes in the IV-plots. G) F-I curves in the first 50 ms immediately after the onset of the current step (*left*) and during a 100-150 ms window of the current step (*right*). Statistical analysis: RM 2W-ANOVA for all panels except M-W t-test for C. *p<0.05, ***p<0.001, ****p<0.0001.

As described for D2 SPNs, we performed two additional analyses of F-I curves to determine the relative roles of subthreshold and suprathreshold changes underlying increased intrinsic excitability in *Foxp1^cKO^* D1 SPNs. First, we replotted the F-I curve data (in figure 6B) at different 50 ms epochs during the current step. Unlike D2 SPNs, we found that the relative differences between *Foxp1^CTL^* and *Foxp1^cKO^* D1 SPNs were similar in both early and later 50 ms epochs (Fig. 4G) suggesting a decreased role for subthreshold membrane properties in the hyperexcitability of *Foxp1^cKO^* D1 SPNs compared to *Foxp1^cKO^* D2 SPNs. (see Fig. 1G).

Next, we replotted F-I curves only for D1 SPNs falling within a limited resting potential range (between - 75 and −90 mV) and found a similar 2-fold increase in the spike numbers (data not shown). However, the average resting membrane potential and input resistance were not detectably different (Table 1). Like D2 SPNs, there was no effect of *Foxp1* deletion on the threshold potential. The threshold current step to evoke a spike was decreased by 17% in *Foxp1^cKO^* D1 SPNs compared to controls (there was a 44% decrease in *Foxp1^cKO^* D2 SPNs, Table 1). Again, these data indicate that subthreshold alterations play a less prominent role in the hyperexcitability of *Foxp1^cKO^* D1 SPNs when compared with *Foxp1^cKO^* D2 SPNs.

### Single cell RNA sequencing data indicate that K_IR_ and K_Leak_ channel transcripts are down-regulated with Foxp1 deletion in D2 SPNs

We analyzed our single-cell RNA-seq (scRNA-seq) dataset(15) from D1 and D2 SPNs to determine whether expression of subthreshold potassium channels are altered when *Foxp1* is deleted. Among K_IR_ channels, KCNJ2 and KCNJ4 are the most highly expressed channels in SPNs (24, 25, 33). Our analysis also showed high expression of their transcripts in *Foxp1^CTL^* D1 and D2 SPNs compared to other *Kcnj* genes (Fig. 5A). However, *Foxp1* deletion significantly reduced their expression in D2 SPNs, consistent with the reduced K_IR_ currents observed in *Foxp1^cKO^* D2 SPNs (Fig. 5B,C). In contrast, only the expression of *Kcnj4* was found to be reduced in *Foxp1^cKO^* D1 SPNs, and this reduction appeared slightly less than that observed in D2 SPNs.

**Figure 5.**
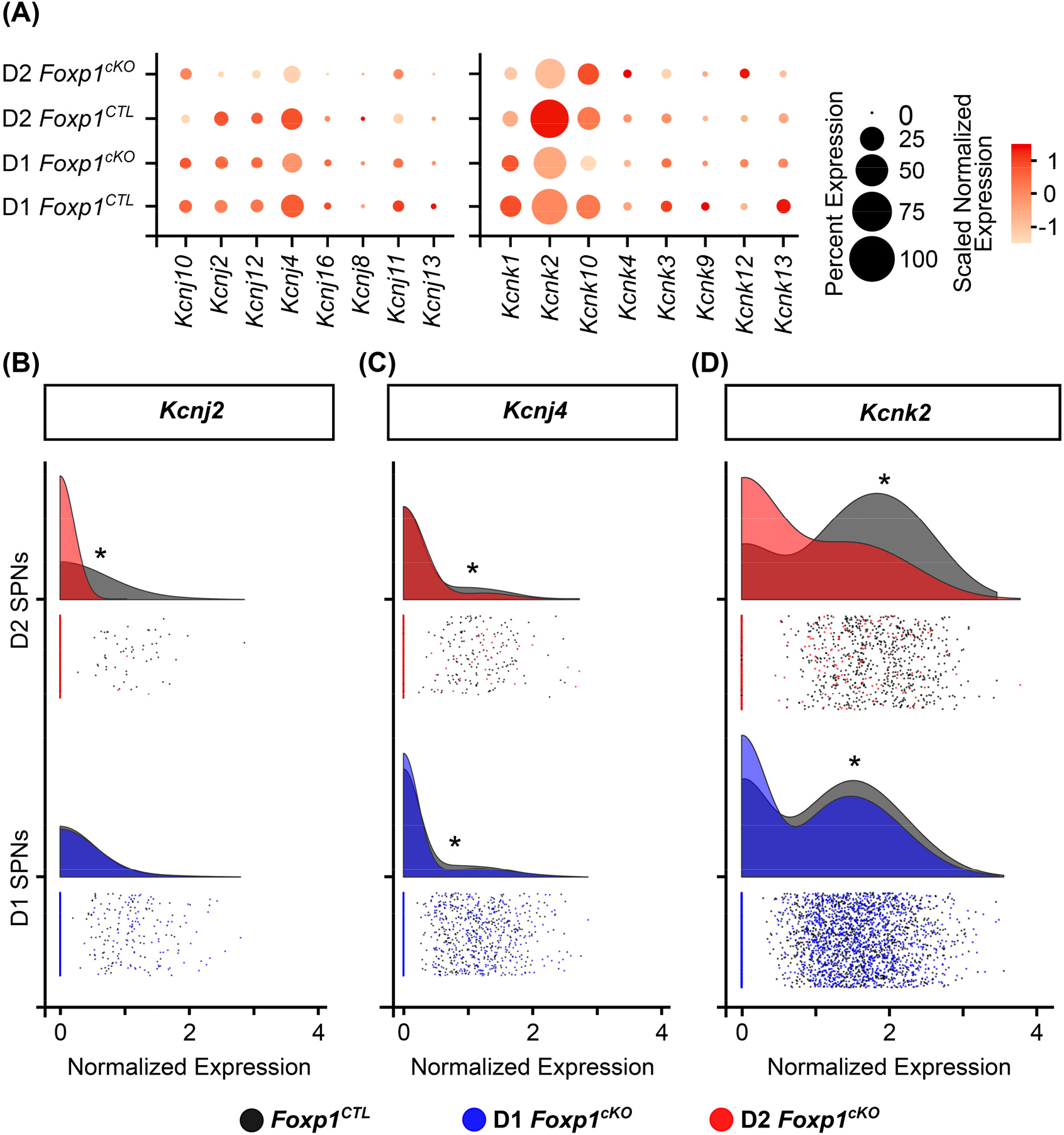
Gene expression changes assayed with single-cell RNA-seq implicate specific K_IR_ and K_Leak_ subtypes being downregulated with *Foxp1* deletion. Gene expression data from our previous study (15) was analyzed for D2 and D1 SPNs with cell-autonomous *Foxp1* deletion using scRNA-seq data. A) Dot plot showing relative expression of transcripts for K_IR_ (*Kcnj*) and K_Leak_ (*Kcnk*) channels across D2 and D1 SPNs. Color of the dots represents scaled log normalized expression of the transcripts whereas size of the dot represents the scaled percentage of SPNs expressing specific transcripts. B-D) RainCloud plot of log normalized expression for *Kcnj2* (B)*, Kcnj4* (C) and *Kcnk2* (D). Cells are grouped as either D2 or D1 SPNs within each genotype and compared across genotypes: *Foxp1^CTL^* compared to D2 *Foxp1^cKO^* for D2 SPNs and *Foxp1C^TL^* compared to D1 *Foxp1^cKO^* for D1 SPNs. In D2 SPNs, all the three transcripts show a significantly down-regulated expression with *Foxp1* deletion (*p<0.05; 0.001, 4.01E-05, 1.32E-101) whereas only *Kcnj4* and *Kcnk2* show a significant reduction in D1 SPNs (*p<0.05; 3.93E-05, 1.69E-92).

The K_Leak_ currents are mediated by the KCNK family of channels. *Kcnk2* is highly expressed in the striatum (28, 29, 31, 32), and consistent with this, we found higher *Kcnk2* transcript expression in *Foxp1^CTL^* D1 and D2 SPNs compared to other *Kcnk* genes (Fig. 5A). Upon *Foxp1* deletion, *Kcnk2* transcript levels were significantly reduced in D2 SPNs (Fig. 5D), suggesting an underlying molecular mechanism for decreased K_Leak_ currents. *Kcnk2* expression was decreased in *Foxp1^cKO^* D1 SPNs as well, however this reduction was less compared to *Foxp1^cKO^* D2 SPNs.

## Discussion

Our study provides the most thorough assessment to date of a role for FOXP1 in the normal physiological development of neurons. We report that FOXP1 negatively regulates intrinsic excitability of D2 SPNs during development through positive regulation of two subthreshold K^+^ currents - K_IR_ and K_Leak_. This regulation is cell-autonomous and likely involves FOXP1 function during both embryonic and postnatal developmental stages. Our pharmacological and single-cell RNA sequencing data indicate that FOXP1 promotes the expression KCNJ2, KCNJ4, and KCNK2 channels, thereby providing specific channel candidates for FOXP1-mediated regulation of K_IR_ and K_Leak_. FOXP1 also appears to negatively regulate intrinsic excitability in D1 SPNs, but unlike D2 SPNs, subthreshold currents play a much smaller role.

### A larger role of K_Leak_ in D2 SPN intrinsic excitability and its regulation by FOXP1

Little is known about the developmental and functional roles of K_Leak_ in SPNs. This partly stems from the lack of pharmacological reagents that can specifically and effectively block the KCNK channels in neurons. Nevertheless, the combination of pharmacological manipulations, *Foxp1* deletion, and single-cell transcriptomics enabled us to examine K_Leak_ in SPNs. In our experiments, reduction in the normalized conductance at −65 to −55 mV membrane potential due to *Foxp1* deletion most likely reflected changes in K_Leak_, and data from previous studies support this assertion. Consistent with Cs^+^ not blocking K_Leak_ (46, 47), we observed minor effects on the conductance of D2 SPNs at these potentials with Cs^+^-application (<10%, Fig. 2C). However, we observed large effects in the presence of Ba^2+^ which is known to block ^K^Leak (41) (Fig. 3A).

We calculated the approximate contribution of a putative K_Leak_ to the total conductance of *Foxp1^CTL^* D2 SPNs as well as the amount that is regulated by *Foxp1* (Table S3). Putative K_Leak_ accounted for 51% of the conductance in *Foxp1^CTL^* D2 SPNs which decreased to 42% in *Foxp1^cKO^* D2 SPNs. This decrease is consistent with our finding of preferential regulation of K_Leak_ by FOXP1 compared to other subthreshold currents (see Fig. 3B,G). Of the down-regulated conductance due to *Foxp1* deletion, K_Leak_ contributed 62%. Even with the possibility that our calculations involve minor contamination by other currents, these calculations suggest a significant impact by K_Leak_ on D2 SPN excitability and its regulation by FOXP1.

### Mechanisms of FOXP1-mediated regulation of K_IR_ and K_Leak_

Our data suggest that FOXP1 negatively regulates the excitability of D2 SPNs through K_IR_ and K_Leak_ by transcriptionally activating their corresponding transcripts - *Kcnj2* and *Kcnj4* for K_IR_ and *Kcnk2* for K_Leak_. Consistent with this, other investigators found that *Kcnk2* and *Kncj4* are up-regulated when human *FOXP1* is overexpressed in mouse striatum (48). Previously, we have shown using ChIP-seq data from human neural progenitor cells that FOXP1 binds to *Kcnj2* and *Kcnk2* promoters (9) (*Kcnk2* regulation is unpublished). Therefore, changes in their expression may involve a direct FOXP1-mediated regulation. Taken together with previous studies, our data indicate that FOXP1-dependent regulation of these channels, either directly or indirectly, may occur at the transcriptional rather than post-transcriptional level. Finally, this involves postnatal processes, and we have made additional observations suggesting that K_IR_ changes emerge earlier in development compared to K_Leak_ changes (see Supplementary Fig. S10).

### Links with alterations in the Foxp1^+/-^ mouse

The *Foxp1*^+/−^ mouse is a good model for *FOXP1*–associated ASD, which is primarily due to heterozygous *FOXP1* mutations. In a previous study, using *Foxp1*^+/−^ mice, we observed changes that closely matched our observations with cell-autonomous homozygous deletion in D2 SPNs - increased intrinsic excitability in D2 SPNs and decreased striatal levels of two of the transcripts described above - *Kcnk2* and *Kcnj2* (9). However, in the current study, hyperexcitability was not observed with cell-autonomous heterozygous *Foxp1* deletion in D2 SPNs (Supplementary Fig. S1). There are two possibilities that may explain this discrepancy: 1) hyperexcitability in *Foxp1*^+/−^ mice is not a cell-autonomous process and involves alterations induced by heterozygosity in other cell types or 2) hyperexcitability is a cell-autonomous process in *Foxp1*^+/−^ mice, but due to delayed deletion in D2 *Foxp1^cHet^* mice (Cre expression at E14-15), this process is not induced. Alternatively, both possibilities could be occurring. In this scenario, because gene dosage is reduced in *Foxp1*^+/−^ mice, the same cell-autonomous mechanism is only induced by additional non-cell-autonomous processes.

While we did not observe any change in the excitability of D1 SPNs in *Foxp1*^+/−^ mice (9) or with postnatal *Foxp1* deletion, these neurons were hyperexcitable in D1 *Foxp1^cKO^* mice. A few reasons may explain this discrepancy. First, FOXP1 reduction in *Foxp1*^+/−^ mice is insufficient to induce hyperexcitability. Second, loss of FOXP1 function may be compensated by FOXP2 (15, 49). Thirdly, D1 *Foxp1^cKO^* mice have significant *Foxp1* deletion in the deep layers of neocortex as well, which include cell populations projecting to SPNs. Therefore, it is possible that hyperexcitability of D1 SPNs in D1 *Foxp1^cKO^* mice is not purely cell-autonomous and is influenced by the homozygous *Foxp1* deletion in cortical neurons. Further experiments are required to resolve this issue.

### The D2 Foxp1^cKO^ mouse model

Although there was a reduction in the number of D2 SPNs in the D2 *Foxp1^cKO^* (see Suppl. Methods), the changes in the excitability and subthreshold conductance were very similar to that observed in *Foxp1*^+/−^ mice (9) and with postnatal *Foxp1* deletion – models where no D2 SPN decrease was observed. Moreover, changes in the relevant K_IR_ and K_Leak_ transcripts were comparable to that observed in *Foxp1*^+/−^ mice (9). These common changes among different deletion strategies indicate that our findings in the D2 *Foxp1^cKO^* mice are relevant to pathological processes occurring in *Foxp1*^+/−^ mice – the model for ASD.

### Conclusion

The functional changes that we describe may be involved in the communication deficits observed both in *FOXP1*-linked ASD patients and in the corresponding *Foxp1*^+/−^ mouse model (2, 9). Moreover, the specific proteins that have been identified as FOXP1 targets could be further studied as potential therapeutic targets for *FOXP1*-linked ASD.

## Acknowledgements

Special thanks to Mathew Harper for technical help. Funding provided by the Simons Foundation (SFARI 573689 and 401220) and NIH (MH102603) to GK and JG.

## Author Contributions

NK, designed and performed experiments, analyzed data, wrote manuscript; SC, performed experiments, analyzed data; VR, designed and performed experiments, analyzed data; AK, analyzed data, created figures; AGA, analyzed data; GK, designed experiments, wrote manuscript; JRG, designed experiments, wrote manuscript.

## Declaration

The authors declare no competing financial interests in relation to the work described.

## References

1. Ferland RJ, Cherry TJ, Preware PO, Morrisey EE, Walsh CA. Characterization of Foxp2 and Foxp1 mRNA and protein in the developing and mature brain. J Comp Neurol. 2003;460(2):266–79.

2. Meerschaut I, Rochefort D, Revencu N, Petre J, Corsello C, Rouleau GA, et al. FOXP1-related intellectual disability syndrome: a recognisable entity. J Med Genet. 2017;54(9):613–23.

3. Siper PM, De Rubeis S, Trelles MDP, Durkin A, Di Marino D, Muratet F, et al. Prospective investigation of FOXP1 syndrome. Mol Autism. 2017;8:57.

4. Satterstrom FK, Kosmicki JA, Wang J, Breen MS, De Rubeis S, An JY, et al. Large-Scale Exome Sequencing Study Implicates Both Developmental and Functional Changes in the Neurobiology of Autism. Cell. 2020;180(3):568–84 e23.

5. Iossifov I, O’Roak BJ, Sanders SJ, Ronemus M, Krumm N, Levy D, et al. The contribution of de novo coding mutations to autism spectrum disorder. Nature. 2014;515(7526):216–21.

6. Sanders SJ, He X, Willsey AJ, Ercan-Sencicek AG, Samocha KE, Cicek AE, et al. Insights into Autism Spectrum Disorder Genomic Architecture and Biology from 71 Risk Loci. Neuron. 2015;87(6):1215–33.

7. Stessman HA, Xiong B, Coe BP, Wang T, Hoekzema K, Fenckova M, et al. Targeted sequencing identifies 91 neurodevelopmental-disorder risk genes with autism and developmental-disability biases. Nat Genet. 2017;49(4):515–26.

8. Bacon C, Schneider M, Le Magueresse C, Froehlich H, Sticht C, Gluch C, et al. Brain-specific Foxp1 deletion impairs neuronal development and causes autistic-like behaviour. Mol Psychiatry. 2015;20(5):632–9.

9. Araujo DJ, Anderson AG, Berto S, Runnels W, Harper M, Ammanuel S, et al. FoxP1 orchestration of ASD-relevant signaling pathways in the striatum. Genes Dev. 2015;29(20):2081–96.

10. Scott-Van Zeeland AA, McNealy K, Wang AT, Sigman M, Bookheimer SY, Dapretto M. No neural evidence of statistical learning during exposure to artificial languages in children with autism spectrum disorders. Biol Psychiatry. 2010;68(4):345–51.

11. Di Martino A, Kelly C, Grzadzinski R, Zuo XN, Mennes M, Mairena MA, et al. Aberrant striatal functional connectivity in children with autism. Biol Psychiatry. 2011;69(9):847–56.

12. Langen M, Leemans A, Johnston P, Ecker C, Daly E, Murphy CM, et al. Fronto-striatal circuitry and inhibitory control in autism: findings from diffusion tensor imaging tractography. Cortex. 2012;48(2):183–93.

13. Radulescu E, Minati L, Ganeshan B, Harrison NA, Gray MA, Beacher FD, et al. Abnormalities in fronto-striatal connectivity within language networks relate to differences in grey-matter heterogeneity in Asperger syndrome. Neuroimage Clin. 2013;2:716–26.

14. Heiman M, Schaefer A, Gong S, Peterson JD, Day M, Ramsey KE, et al. A translational profiling approach for the molecular characterization of CNS cell types. Cell. 2008;135(4):738–48.

15. Anderson AG, Kulkarni A, Harper M, Konopka G. Single-Cell Analysis of Foxp1-Driven Mechanisms Essential for Striatal Development. Cell Rep. 2020;30(9):3051–66 e7.

16. Precious SV, Kelly CM, Reddington AE, Vinh NN, Stickland RC, Pekarik V, et al. FoxP1 marks medium spiny neurons from precursors to maturity and is required for their differentiation. Exp Neurol. 2016;282:9–18.

17. Fong WL, Kuo HY, Wu HL, Chen SY, Liu FC. Differential and Overlapping Pattern of Foxp1 and Foxp2 Expression in the Striatum of Adult Mouse Brain. Neuroscience. 2018;388:214–23.

18. Usui N, Araujo DJ, Kulkarni A, Co M, Ellegood J, Harper M, et al. Foxp1 regulation of neonatal vocalizations via cortical development. Genes Dev. 2017;31(20):2039–55.

19. Dasen JS, De Camilli A, Wang B, Tucker PW, Jessell TM. Hox repertoires for motor neuron diversity and connectivity gated by a single accessory factor, FoxP1. Cell. 2008;134(2):304–16.

20. Rousso DL, Gaber ZB, Wellik D, Morrisey EE, Novitch BG. Coordinated actions of the forkhead protein Foxp1 and Hox proteins in the columnar organization of spinal motor neurons. Neuron. 2008;59(2):226–40.

21. Palmesino E, Rousso DL, Kao TJ, Klar A, Laufer E, Uemura O, et al. Foxp1 and lhx1 coordinate motor neuron migration with axon trajectory choice by gating Reelin signalling. PLoS Biol. 2010;8(8):e1000446.

22. Li X, Han X, Tu X, Zhu D, Feng Y, Jiang T, et al. An Autism-Related, Nonsense Foxp1 Mutant Induces Autophagy and Delays Radial Migration of the Cortical Neurons. Cereb Cortex. 2019;29(7):3193–208.

23. Araujo DJ, Toriumi K, Escamilla CO, Kulkarni A, Anderson AG, Harper M, et al. Foxp1 in Forebrain Pyramidal Neurons Controls Gene Expression Required for Spatial Learning and Synaptic Plasticity. J Neurosci. 2017;37(45):10917–31.

24. Shen W, Tian X, Day M, Ulrich S, Tkatch T, Nathanson NM, et al. Cholinergic modulation of Kir2 channels selectively elevates dendritic excitability in striatopallidal neurons. Nat Neurosci. 2007;10(11):1458–66.

25. Cazorla M, Shegda M, Ramesh B, Harrison NL, Kellendonk C. Striatal D2 receptors regulate dendritic morphology of medium spiny neurons via Kir2 channels. J Neurosci. 2012;32(7):2398–409.

26. Nisenbaum ES, Wilson CJ. Potassium currents responsible for inward and outward rectification in rat neostriatal spiny projection neurons. J Neurosci. 1995;15(6):4449–63.

27. Lieberman OJ, McGuirt AF, Mosharov EV, Pigulevskiy I, Hobson BD, Choi S, et al. Dopamine Triggers the Maturation of Striatal Spiny Projection Neuron Excitability during a Critical Period. Neuron. 2018;99(3):540–54 e4.

28. Fink M, Duprat F, Lesage F, Reyes R, Romey G, Heurteaux C, et al. Cloning, functional expression and brain localization of a novel unconventional outward rectifier K+ channel. EMBO J. 1996;15(24):6854–62.

29. Talley EM, Solorzano G, Lei Q, Kim D, Bayliss DA. Cns distribution of members of the two-pore-domain (KCNK) potassium channel family. J Neurosci. 2001;21(19):7491–505.

30. Feliciangeli S, Chatelain FC, Bichet D, Lesage F. The family of K2P channels: salient structural and functional properties. The Journal of physiology. 2015;593(12):2587–603.

31. Heurteaux C, Guy N, Laigle C, Blondeau N, Duprat F, Mazzuca M, et al. TREK-1, a K+ channel involved in neuroprotection and general anesthesia. EMBO J. 2004;23(13):2684–95.

32. Maze I, Chaudhury D, Dietz DM, Von Schimmelmann M, Kennedy PJ, Lobo MK, et al. G9a influences neuronal subtype specification in striatum. Nat Neurosci. 2014;17(4):533–9.

33. Karschin C, Dissmann E, Stuhmer W, Karschin A. IRK(1-3) and GIRK(1-4) inwardly rectifying K+ channel mRNAs are differentially expressed in the adult rat brain. J Neurosci. 1996;16(11):3559–70.

34. Lieberman OJ, Frier MD, McGuirt AF, Griffey CJ, Rafikian E, Yang M, et al. Cell-type specific regulation of neuronal intrinsic excitability by macroautophagy. Elife. 2020;9.

35. Simons Foundation. Gene Scoring New York, NY: Simons Foundation; 2020

36. Harrington AJ, Raissi A, Rajkovich K, Berto S, Kumar J, Molinaro G, et al. MEF2C regulates cortical inhibitory and excitatory synapses and behaviors relevant to neurodevelopmental disorders. Elife. 2016;5.

37. Kahanovitch U, Cuddapah VA, Pacheco NL, Holt LM, Mulkey DK, Percy AK, et al. MeCP2 Deficiency Leads to Loss of Glial Kir4.1. eNeuro. 2018;5(1).

38. Mermelstein PG, Song WJ, Tkatch T, Yan Z, Surmeier DJ. Inwardly rectifying potassium (IRK) currents are correlated with IRK subunit expression in rat nucleus accumbens medium spiny neurons. J Neurosci. 1998;18(17):6650–61.

39. Hagiwara S, Miyazaki S, Rosenthal NP. Potassium current and the effect of cesium on this current during anomalous rectification of the egg cell membrane of a starfish. J Gen Physiol. 1976;67(6):621–38.

40. Hibino H, Inanobe A, Furutani K, Murakami S, Findlay I, Kurachi Y. Inwardly rectifying potassium channels: their structure, function, and physiological roles. Physiol Rev. 2010;90(1):291–366.

41. Ma XY, Yu JM, Zhang SZ, Liu XY, Wu BH, Wei XL, et al. External Ba2+ block of the two-pore domain potassium channel TREK-1 defines conformational transition in its selectivity filter. J Biol Chem. 2011;286(46):39813–22.

42. Sandoz G, Douguet D, Chatelain F, Lazdunski M, Lesage F. Extracellular acidification exerts opposite actions on TREK1 and TREK2 potassium channels via a single conserved histidine residue. Proc Natl Acad Sci U S A. 2009;106(34):14628–33.

43. Mazella J, Petrault O, Lucas G, Deval E, Beraud-Dufour S, Gandin C, et al. Spadin, a sortilin-derived peptide, targeting rodent TREK-1 channels: a new concept in the antidepressant drug design. PLoS Biol. 2010;8(4):e1000355.

44. Djillani A, Pietri M, Mazella J, Heurteaux C, Borsotto M. Fighting against depression with TREK-1 blockers: Past and future. A focus on spadin. Pharmacol Ther. 2019;194:185–98.

45. Djillani A, Pietri M, Moreno S, Heurteaux C, Mazella J, Borsotto M. Shortened Spadin Analogs Display Better TREK-1 Inhibition, In Vivo Stability and Antidepressant Activity. Front Pharmacol. 2017;8:643.

46. Lesage F, Lazdunski M. Molecular and functional properties of two-pore-domain potassium channels. Am J Physiol Renal Physiol. 2000;279(5):F793–801.

47. Coetzee WA, Amarillo Y, Chiu J, Chow A, Lau D, McCormack T, et al. Molecular diversity of K+ channels. Ann N Y Acad Sci. 1999;868:233–85.

48. Tang B, Becanovic K, Desplats PA, Spencer B, Hill AM, Connolly C, et al. Forkhead box protein p1 is a transcriptional repressor of immune signaling in the CNS: implications for transcriptional dysregulation in Huntington disease. Hum Mol Genet. 2012;21(14):3097–111.

49. Vernes SC, Oliver PL, Spiteri E, Lockstone HE, Puliyadi R, Taylor JM, et al. Foxp2 regulates gene networks implicated in neurite outgrowth in the developing brain. PLoS Genet. 2011;7(7):e1002145.

