## Supplementary figures and images for "FOXP1 negatively regulates intrinsic excitability in D2 striatal projection neurons by promoting inwardly rectifying and leak potassium currents"

### Supplementary Fig 1

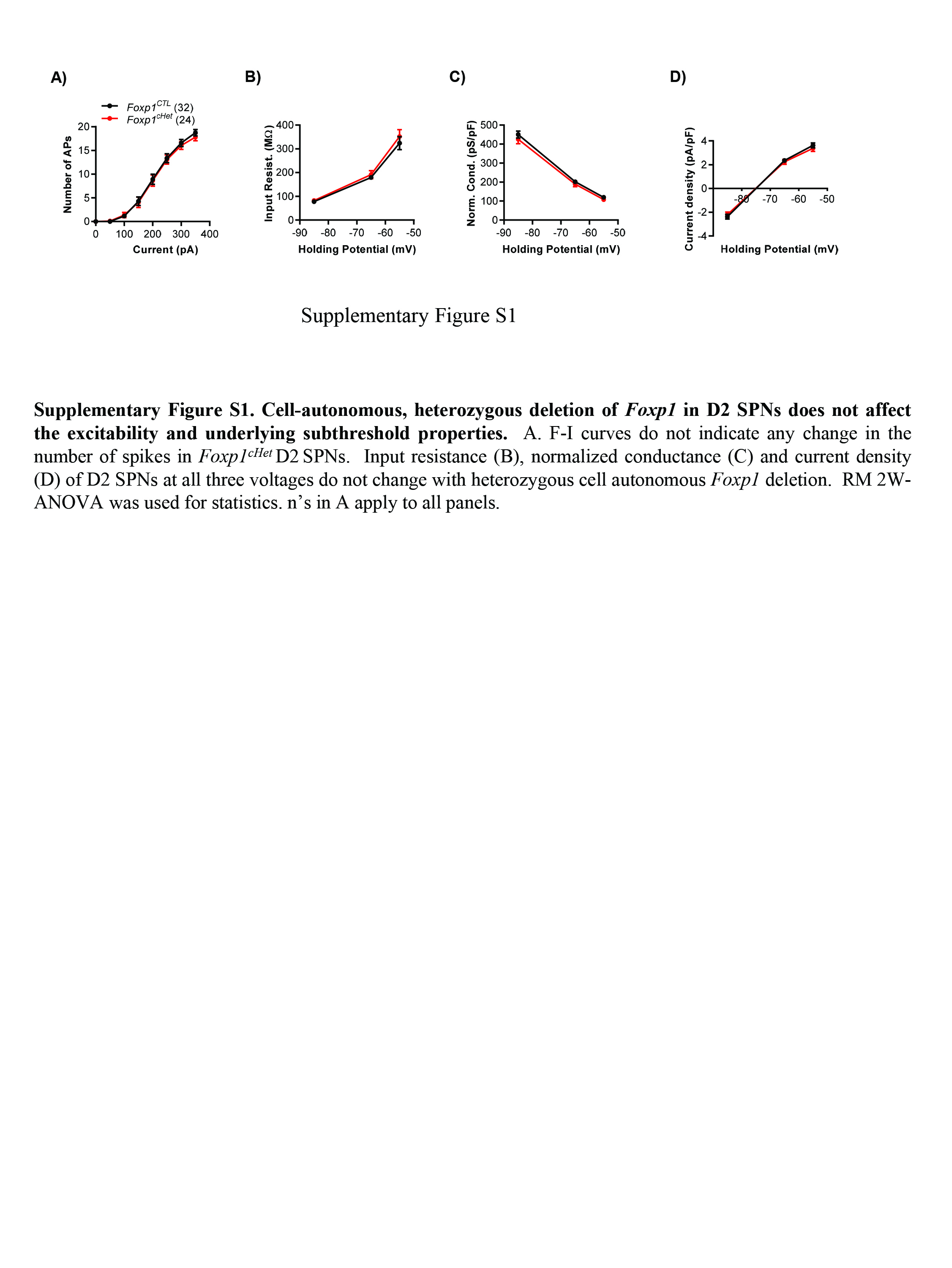

### Supplementary Fig 2

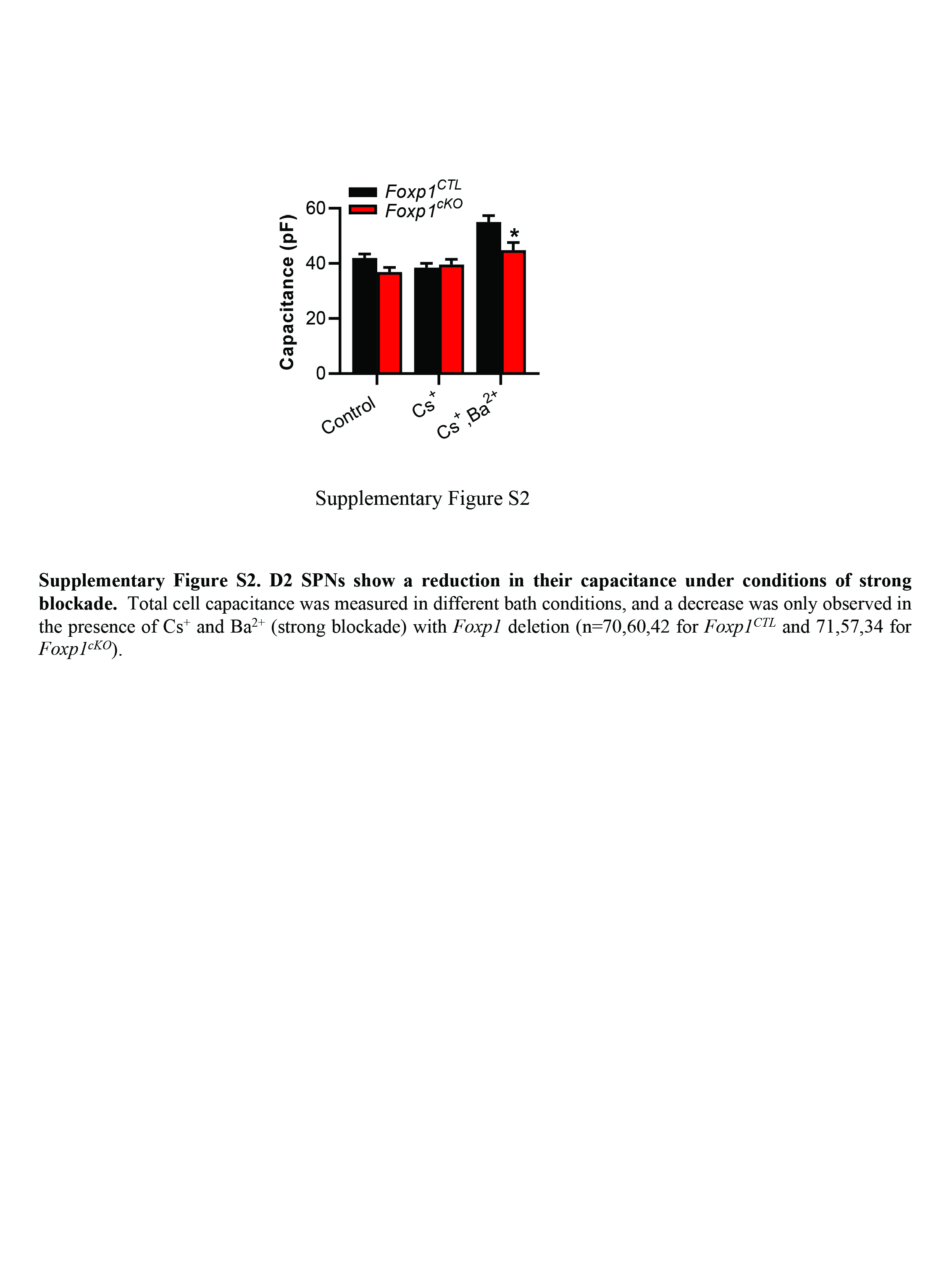

### Supplementary Fig 3

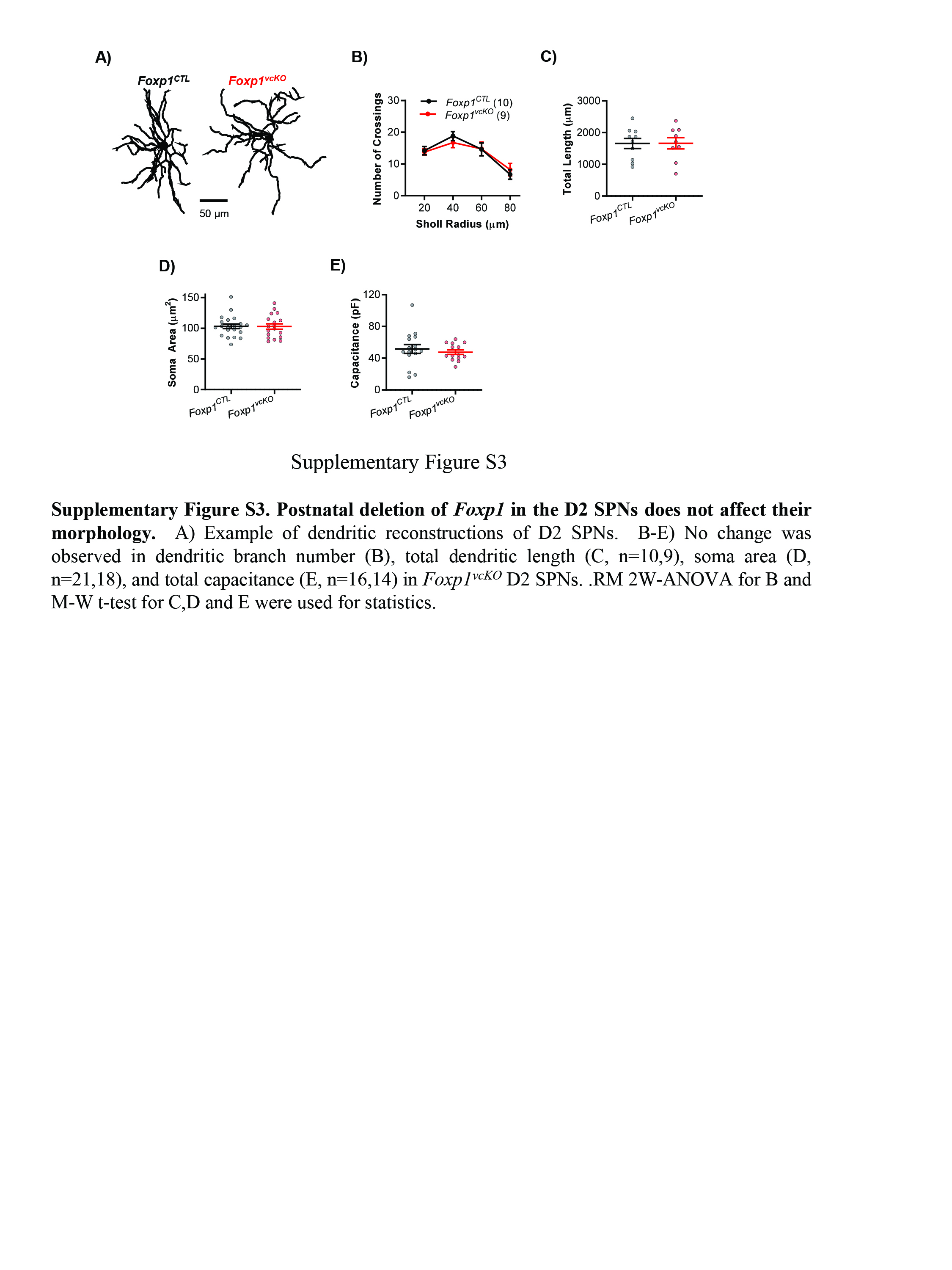

### Supplementary Fig 4

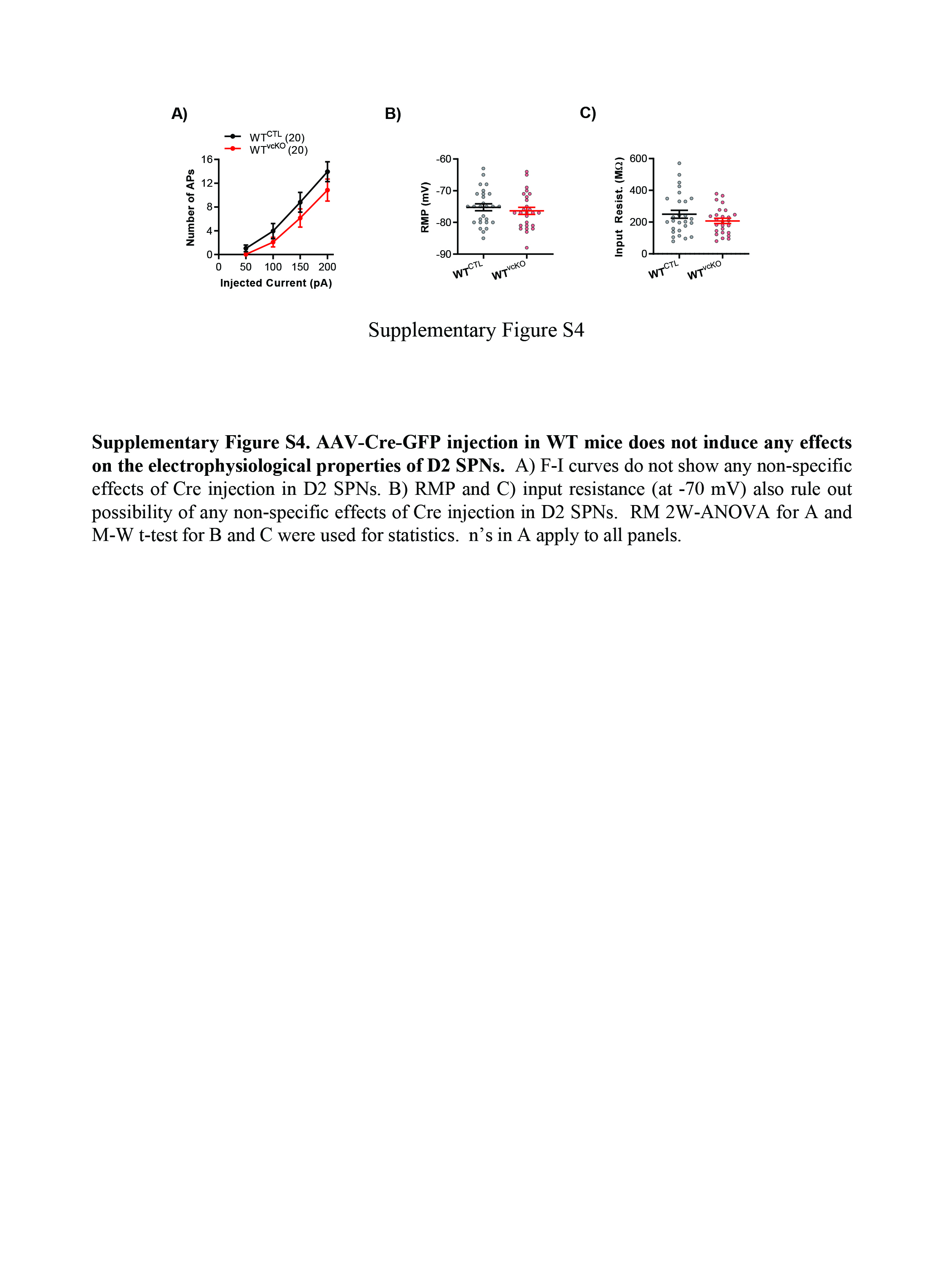

### Supplementary Fig 5

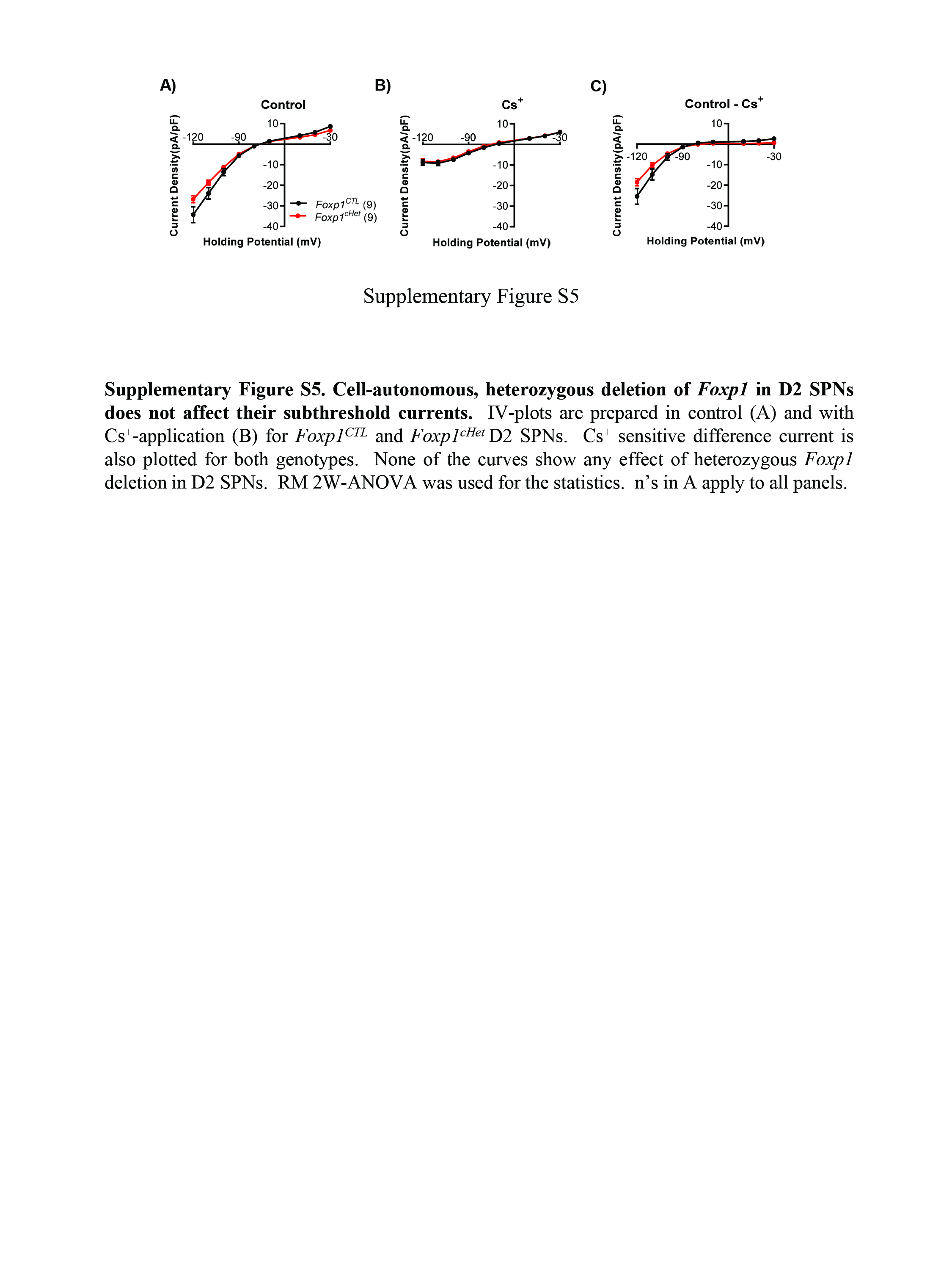

### Supplementary Fig 6

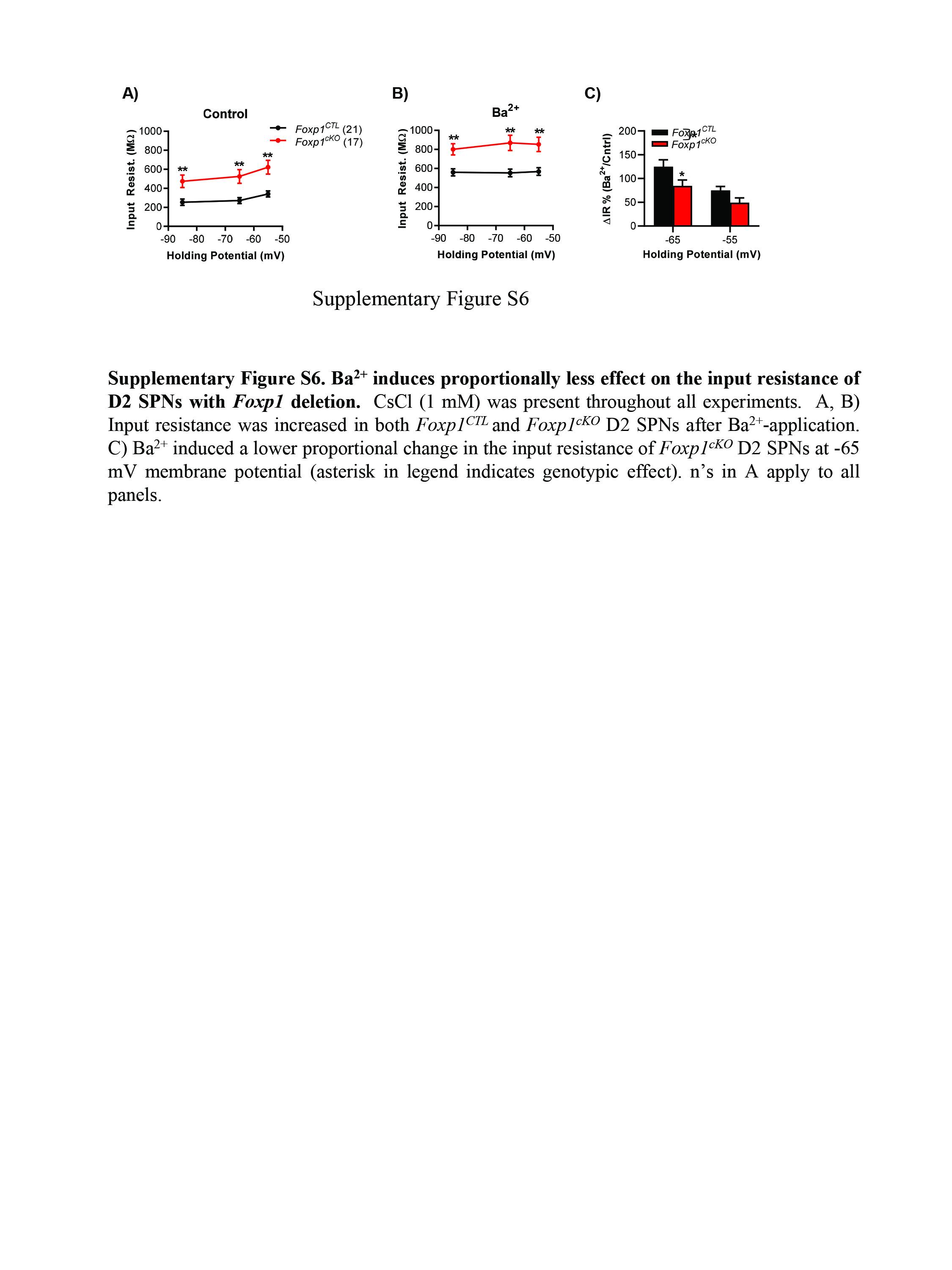

### Supplementary Fig 7

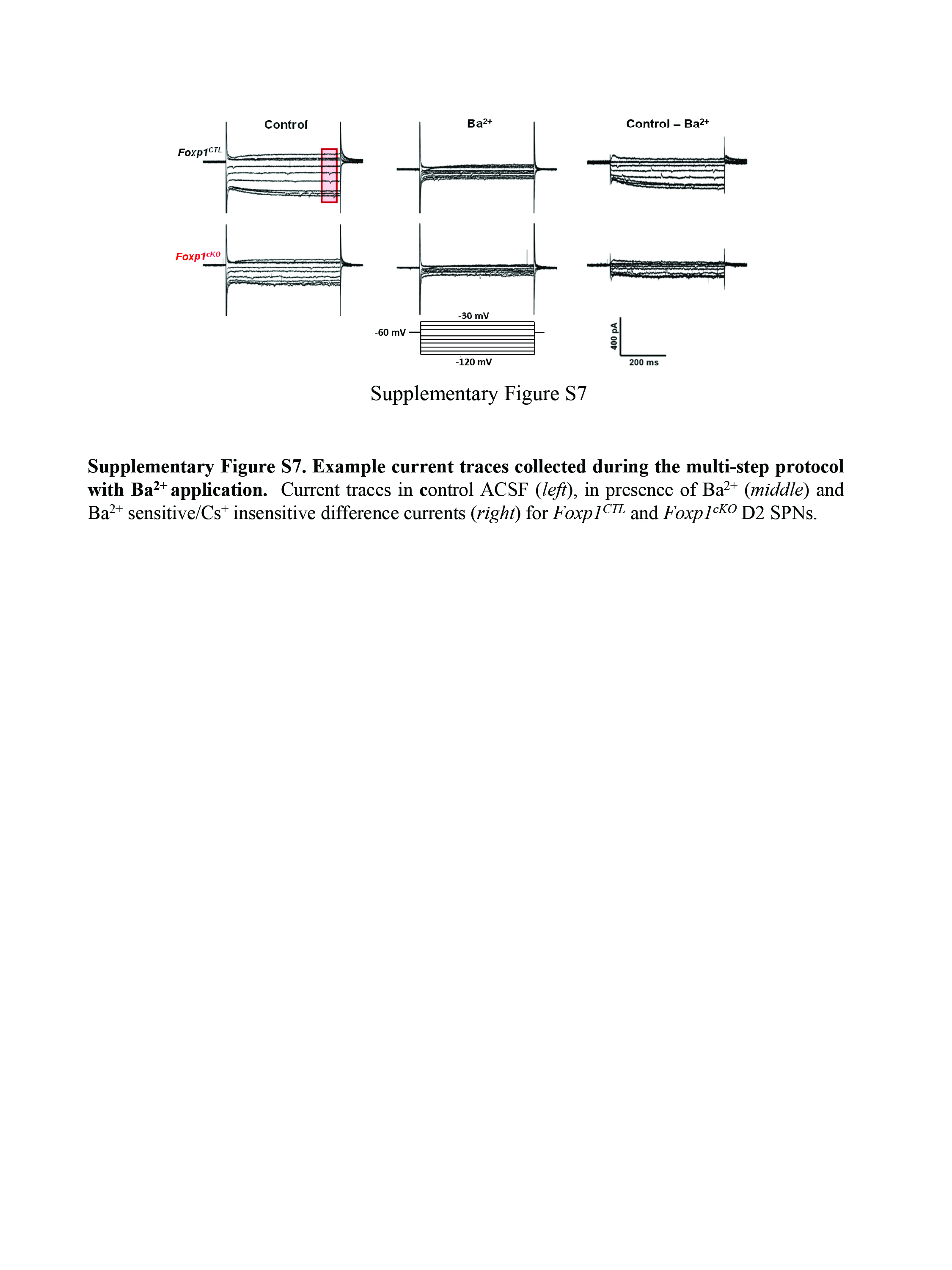

### Supplementary Fig 8

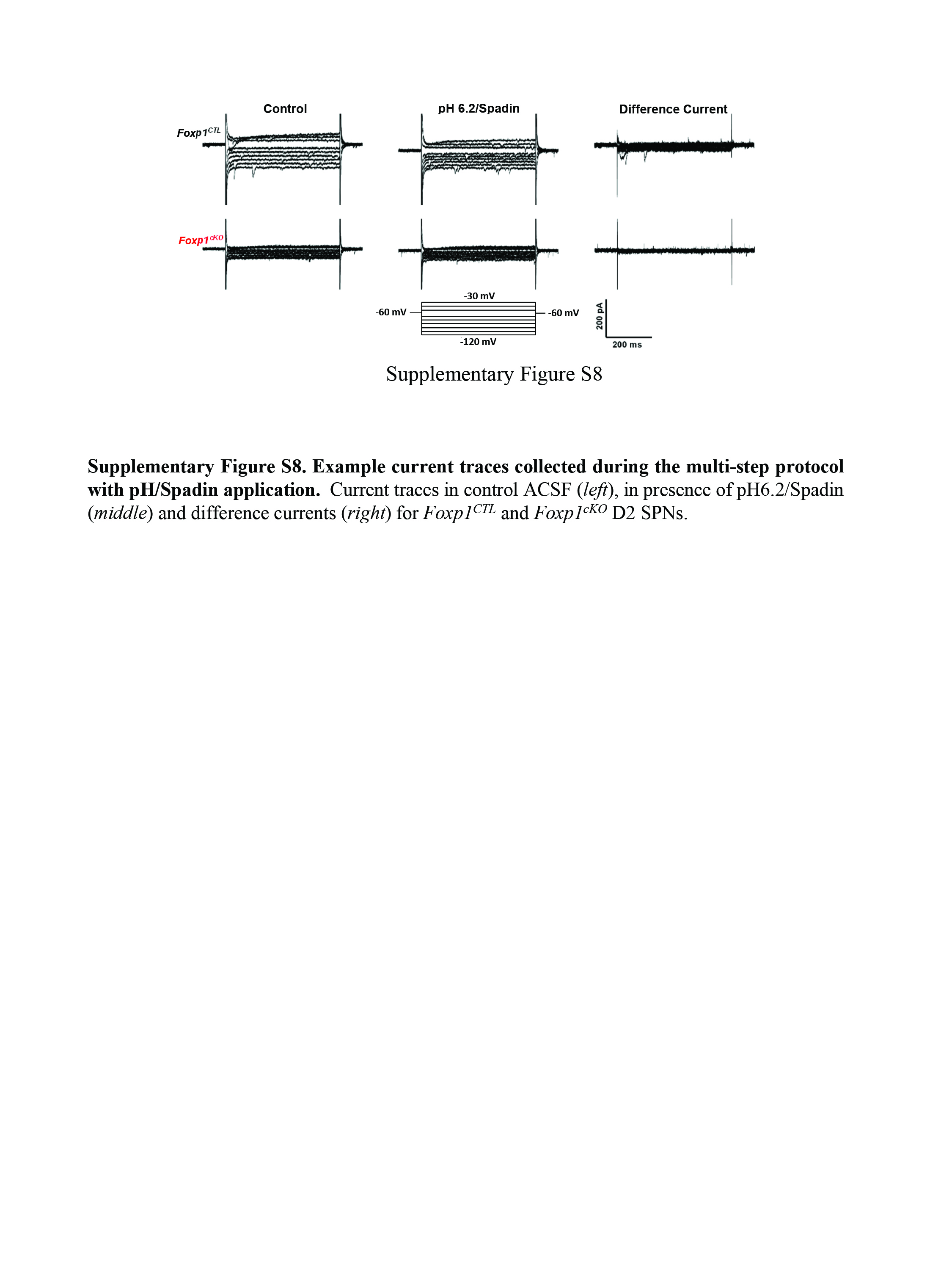

### Supplementary Fig 9

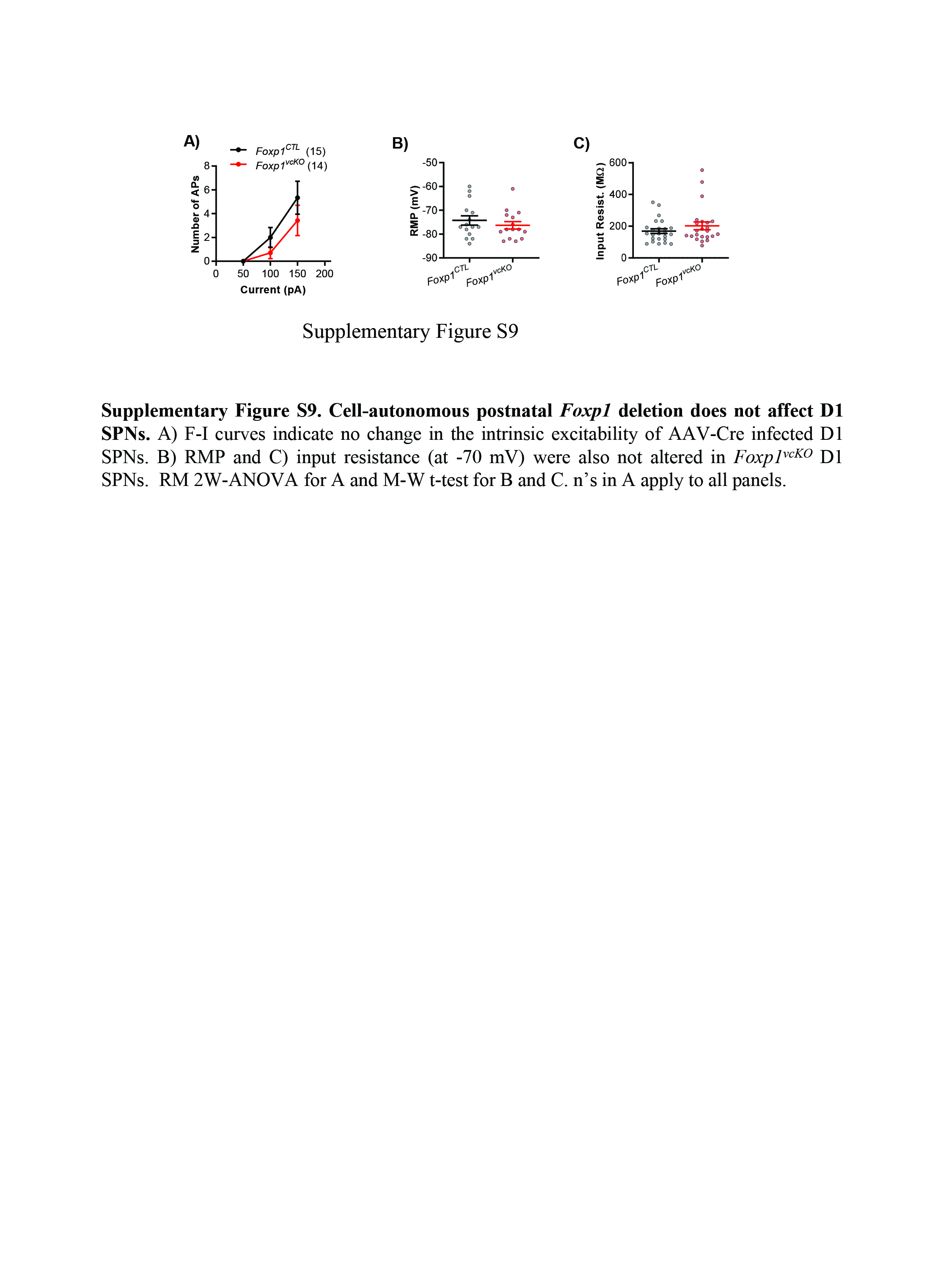

### Supplementary Fig 10

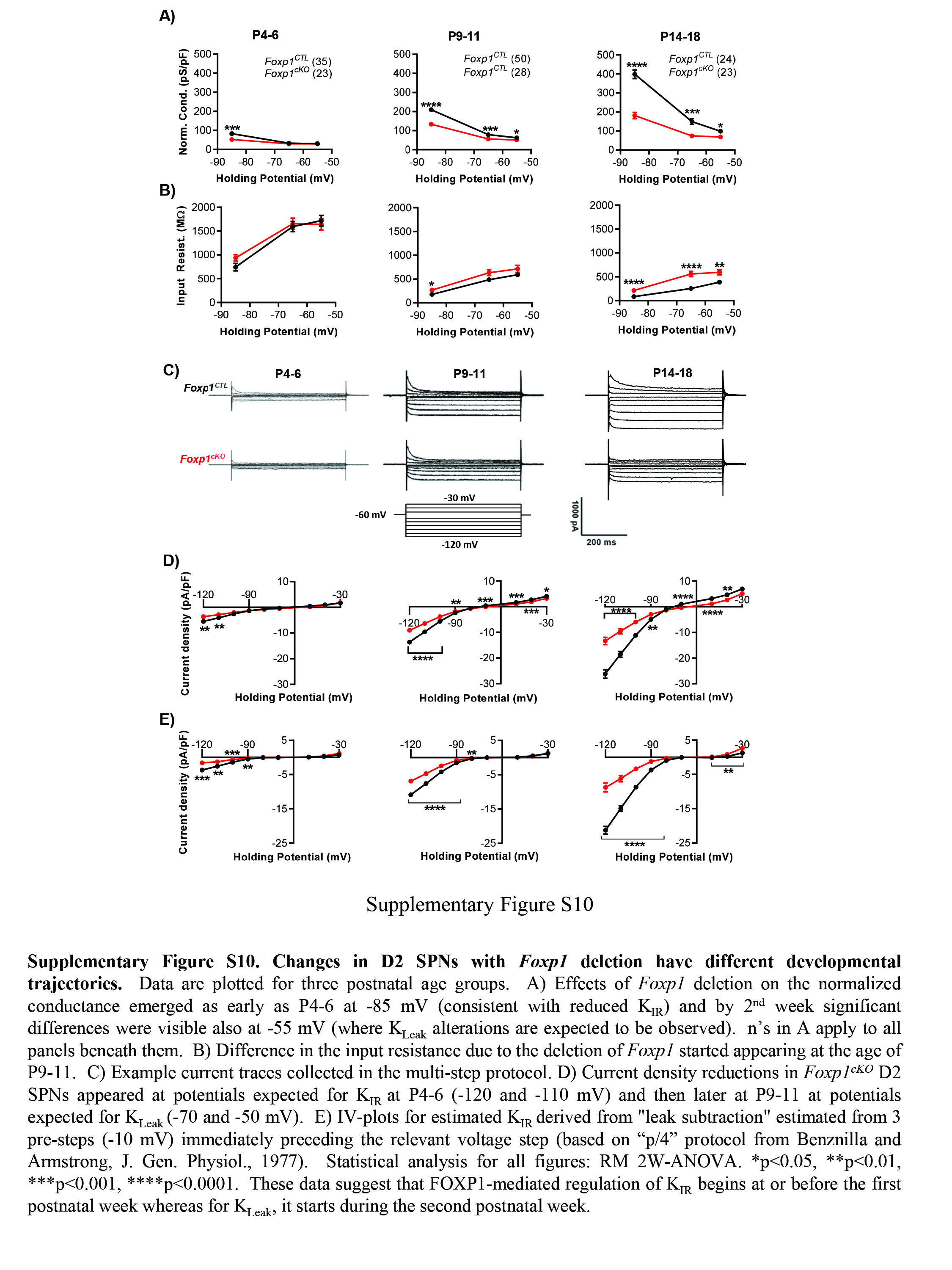
