## Supplementary Methods for "FOXP1 negatively regulates intrinsic excitability in D2 striatal projection neurons by promoting inwardly rectifying and leak potassium currents"

### *Mice*

All mice used were of C57BL/6J background strain. *Foxp1*$^{flox/flox}$ mice were provided by Dr. Haley Tucker. These mice were further backcrossed to C57BL/6J for at least 10 generations to obtain congenic animals as previously described [1,2]. *Drd2*-Cre$^{tg/-}$ and *Drd1α*-Cre$^{tg/-}$ mice were BAC-transgenic mice driving Cre expression under the promoters of *Drd2-receptor* (ER44Gsat, 032108-UCD) and *Drd1α-receptor* (262Gsat, 030989-UCD), respectively [3]. We also used two BAC-transgenic fluorescent mouse strains: *Drd2*-eGFP$^{tg/-}$ and *Drd1α*-tdTomato$^{tg/-}$ [3,4].

*Drd2* conditional *Foxp1* knockout (D2 *Foxp1*$^{cKO}$) mice were transgenic for three alleles: *Drd2*-Cre$^{tg/-}$:*Foxp1*$^{flox/flox}$:*Drd2*-eGFP$^{tg/-}$. Littermates of this group that did not express Cre served as WT controls (*Foxp1*$^{CTL}$). The analogous description applies for *Drd1α Foxp1*$^{cKO}$ (D1 *Foxp1*$^{cKO}$) mice which were also triple transgenic: *Drd1α*-Cre$^{tg/-}$:*Foxp1*$^{flox/flox}$:*Drd1α*-tdTomato$^{tg/-}$. We also employed a heterozygous version of D2 *Foxp1*$^{cKO}$ which are referred to by D2 *Foxp1*$^{cHet}$. D2 *Foxp1*$^{cKO}$ mice have a smaller striatum and approximately one-third of the normal D2-SPN population [5]. The remaining D2 SPNs in D2 *Foxp1*$^{cKO}$ mice were healthy as they showed stable resting potentials and fired action potentials during our recordings. In D1 *Foxp1*$^{cKO}$ mice, the number of D2 SPNs was unaltered. Based on a percentage measurement of different striatal cell types, D1 SPN number did not appear decreased in either *Foxp1*$^{cKO}$ mouse model, but this was not directly measured.

Both D2 and D1 *Foxp1*$^{cKO}$ mice express Cre which co-localize with FOXP1 in the striatum between by embryonic day 14-15 (E14-15) based on experiments performed using a Cre-dependent Rosa-YFP mice [5]. *Foxp1* expression turns on earlier at E12, and in both *Foxp1*$^{cKO}$ mouse lines, both protein and mRNA levels are maximally reduced in the striatum by P7 [5]. Both male and female mice were used for experiments. Mice were maintained on a 12 h light on/off schedule with food and water *ab limitum*. All experiments were performed according to the procedures approved by the UT Southwestern Institutional Animal Care and Use Committee.

### *Viral-mediated postnatal deletion of Foxp1*

*Foxp1* was deleted postnatally using AAV-mediated Cre expression. AAV9 viral particles for the expression of eGFP-Cre fusion protein (pAAV-CMV-HI.eGFP-Cre) were obtained from the University of Pennsylvania Vector Core, Gene Therapy Program. The viral particles were diluted to $10^{12}$ pfu in sterile 1X PBS and injections were performed as described previously [6,7]. In brief, 1 day old *Foxp1*$^{flox/flox}$:*Drd1α*-tdTomato$^{tg/-}$ C57BL/6J pups were anesthetized on ice, and 350 nl of viral particles were injected into the left striatum at a depth of 1.5 mm using a beveled glass injection pipette and a Nanoject™ injector (Drummond Scientific, Inc.). Pups were recovered on a heating pad until they regained consciousness and then transferred to their home cage. Mice positive for the tdTomato reporter were used for experiments. Under the fluorescence microscope, D1 SPNs expressed tdTomato, and D2 SPNs were tdTomato-negative. This identification of D2 SPNs was reliable based on the smaller size of SPNs and the fact that 95% of striatal neurons are SPNs [4]. At the same time, eGFP fluorescence in nuclei marked Cre expression through viral infection. Therefore, expression of tdTomato and eGFP permitted identification of the SPN subtypes and expression of *Foxp1*. Approximately 30% of SPNs, on average, were detectably infected as observed by GFP fluorescence in their nuclei. Previous studies report Cre-expression and deletion by P5-P10 using this virus [6,7].

*Brain slices and Recordings*

Unless stated otherwise, postnatal day (P) 14-23 mice were used for the electrophysiological recordings. Mice older than P9 were anesthetized with a mixture of ketamine (125 mg/kg) and xylazine (25 mg/kg), and brains were quickly dissected and immersed in partially frozen dissection buffer [8]. Neonatal mice P4-6 were anesthetized by hypothermia by keeping on ice for 5-10 min and then decapitated to dissect their brain. Using a Vibratome (Leica VTS 1200S), 300-350 µm (400 µm for P4-6) thick brain slices containing the dorsal striatum were cut at ~4°C in dissection buffer. These slices were immediately transferred to nominal ACSF (bubbled with 5% $CO_2$ and 95% $O_2$) first at 35°C for 30 min, and then at room temperature for a minimum of 30 min before recordings [8,9]. D1 or D2 SPNs from the dorsal striatum were targeted based on fluorescent reporter expression for whole-cell recording using glass pipettes of 5-8 MΩ resistance. Only neurons with a maximum acceptable resting membrane potential (-50 mV for >P9, -45 mV for P4-6), capacitance greater than 15 pF, stable baseline current within 15 pA, and series resistance less than 20 MΩ were analyzed. For all experiments, average series resistance did not differ between *Foxp1*$^{cKO}$ and *Foxp1*$^{CTL}$ neurons. The junction potential was ~10 mV and was not corrected unless stated otherwise. Therefore, all membrane potentials are approximately in error by +10 mV.

For slices obtained from *Foxp1*$^{cKO}$ mice, recordings were performed at 30ºC. Only D2 SPNs were recorded in D2 *Foxp1*$^{cKO}$ mice, where WT controls and *Foxp1*-deleted neurons were named *Foxp1*$^{CTL}$ and *Foxp1*$^{cKO}$ D2 SPNs, respectively. Similar nomenclature applies to D1 SPNs. For the slices where *Foxp1* was deleted postnatally using AAV-mediated Cre expression, recording temperature was kept at 26ºC, and uninfected SPNs were named *Foxp1*$^{CTL}$ SPNs while AAV-infected neurons were named *Foxp1*$^{vcKO}$ SPNs.

*Current steps, action potential shape, and analysis*

In current clamp, incremental current steps (500 ms duration) were applied at resting potential to measure intrinsic excitability. The number of action potentials elicited at each step are counted and used to plot a "firing versus current injection" curve, or "F-I" curve.

Action potentials (spikes) obtained during the current steps are analyzed for several parameters. The threshold current is the minimum current step required to evoke an action potential. Threshold potential is calculated at the initial upward inflection of the spike defined by voltage at which the 3$^{rd}$ derivative of the trace peaks. This is done for the spikes evoked at the threshold current. We also perform an "equalized resting-potential potential (RMP)" analysis of spiking properties by removing neurons that do not fall within a given resting membrane potential range (For D1 and D2 neurons, approximately 15% of neurons were removed, see Table 1). These criteria make conditions for quantifying spiking more equalized and less affected by depolarization-induced currents.

We also perform waveform analysis on the spikes evoked by the current steps by measuring their height, width, and latency (Table 1). One trace is selected from each cell where a current step resulted in at least 3 spikes with a firing rate of 8 Hz (±6 Hz). Spike height is the difference between the initial inflection point (defined by a peak in the 3$^{rd}$ derivative of the trace) and the spike peak. Spike width is measured at the halfway point of the spike height. Finally, we measure the latency to the first spike relative to the current step onset. We have presented the measurements for the first spike waveform only, however the pattern was consistent among the 3 spikes.

*Single -10 voltage step and analysis to measure input resistance, capacitance, and conductance*

In voltage clamp, a single -10 mV voltage step (500 ms duration) was applied to measure input resistance, capacitance, normalized cell conductance, and the voltage-dependence of holding current. Input resistance and conductance are calculated based on average current values in a 100 ms window before and starting at 200 ms after step onset. Capacitance is calculated by measuring the width at half-height ($W_{HH}$) of the transient induced immediately after the voltage step (as an approximation of the decay time constant), and then using the equation, $C = W_{HH} / R_S$, where C is capacitance and $R_S$ is series resistance [10]. Normalized conductance is calculated by dividing the conductance by the capacitance.

For recordings obtained from the *Foxp1$^{cKO}$* mice, the -10 steps were applied at -85, -65, and -55 mV holding potentials. However, for experiments where *Foxp1* was deleted postnatally using AAV-mediated Cre expression, this was done only at -70 mV holding potential.

*Multi-step protocol using IV-plots*

In voltage clamp, a multi-step protocol (500 ms duration) was applied while maintaining a -60 mV holding potential (ranging from -30 to -120 mV voltages in -10 mV steps). Series resistance compensation was applied (bandwidth = 1.02 KHz, correction and prediction = 50%). The current measured was the average of a 200 ms window at the end of the step – a time where steady state current was attained (see Fig. 2E). Three -10 mV steps of the same duration preceded the above voltage steps so that we could perform linear subtraction of estimated passive currents, but while we examined this, we do not report these data. Current values were divided by the capacitance to obtain current density, however for the ease of explanation, at several places we have used the term 'current' which always refers to 'current density' only.

We make a "genotypic" comparison of average IV-plots between $Foxp1^{CTL}$ and $Foxp1^{cKO}$ D2 SPNs using a 2-Way ANOVA (genotype x voltage step) with repeated measures for the voltage steps. This is followed by post-hoc multi-comparison tests. Genotypic *difference* IV-plots are generated by subtracting average $Foxp1^{cKO}$ plots from average $Foxp1^{CTL}$ plots. Standard error for each step is calculated by taking the square root of the sum squared SE values of the respective currents [11,12].

We also make a "reagent" comparison of average IV-plots between "before" control and "after" reagent application for a single genotype using a 2-Way ANOVA (reagent x voltage step) with repeated measures for both variables. This is followed by post-hoc multi-comparison tests. For each neuron, currents in the presence of reagent are subtracted from those obtained before its application (control conditions). These difference currents are averaged for both genotypes to obtain reagent *difference* IV-plots.

*Membrane potentials for measuring $K_{IR}$ and $K_{Leak}$*

We focus on two membrane potential ranges for the analysis of $K_{IR}$ and $K_{Leak}$ currents (briefly referred as $K_{IR}$ and $K_{Leak}$). The first range is defined by membrane potentials where we observed $K_{IR}$ (from -120 mV through -85 mV). The rationale for this is supported by the $K_{IR}$ IV-plots in figure 2F. The second range is define by the -70 to -50 mV range where the results in our experiments are most likely attributed to $K_{Leak}$ since: 1) most of the currents activated at these potentials are blocked by the compounds in our cocktail (Supplementary Table S1, S2); 2) most of the depolarization-activated currents are not significantly activated (with junction potential taken into account, -50 mV is actually ~-60 mV), 3) current is measured in the later part of the step (steady state), and hence, any transient voltage-dependent current occurring at the beginning of the step was avoided; 4) the driving force for $K_{Leak}$ is high, 5) the conductance of $K_{Leak}$ channels is higher compared to more hyperpolarized potentials due to their outward rectification [13], and 6) $K_{IR}$ is undetectable at these potentials (see Fig. 2F) and is mostly blocked in the experiments where we measure $K_{Leak}$ (Fig. 3). Previous studies have also used this range to observe $K_{Leak}$ [12,14]. For experiments in figure 3 (*Right*) when $Ba^{2+}$ was continually present, conductance results at all membrane potentials were in the context of $K_{Leak}$ because of the thorough blockade of all other currents. For IV-plots in figure 3 (*Right*), we focused on membrane potentials at and above -70 mV to examine $K_{Leak}$ because of the greater driving force.

Because KCNQ ($K_v$7.1-7.5) channels are open at -60 to -50 mV, they could confound our assumption of collecting data relevant to $K_{Leak}$ at these potentials. But 3 reasons make this possible confound unlikely: 1) TEA-Cl in the bath blocks 50%-80% of these currents [15,16] (Supplementary Table S1), 2) KCNQ channels are minimally open at these potentials [15,16], and 3) The replacement of cytosolic solution with pipette solution during whole-cell recording inactivates KCNQ channels [15,17].

*Morphology and analysis*

Dendritic morphology was measured by filling neurons with a fluorescent dye (AF488, 100 µM, Life Technologies, Inc.) and biocytin (2 mg/ml, Life Technologies, Inc.) during the whole-cell recording. The former was used for confirmation of the dye diffusion during recording and the later was used for post-fixation measurements. After recording, cells in the slices were fixed in 2% PFA/4% sucrose solution for 30 min at room temperature and then for 18-24 h at 4ºC, which also quenched AF488 filling dye. After fixation, slices were washed three times with 1X phosphate buffer saline (PBS) followed by 1 h of blocking and permeabilization (10% NDS/ 0.6% Triton-X-100 in 0.1 M PBS). Thereafter, slices were incubated with streptavidin-conjugated AF488 or AF555 (10 µl/ml) for 40 min. Slices were again washed three times and then mounted for imaging on an LSM 510 inverted confocal fluorescence microscope, and Z-stack images were acquired. The dendritic arbor of each filled cell was traced through the Z-stack using the Simple Neurite Tracer plug-in (Fiji/Image-J) [18,19]. Sholl analysis is performed using concentric radii step intervals of 20 µm and dendritic crossings at each radius are counted. Two-way ANOVA with repeated measures is applied for the statistical measurements.

*Electrophysiology Solutions*

Brain slices were cut in dissection buffer with the following components (mM): 110 Choline-Cl, 2.5 KCl, 1.25 $NaH_2PO_4$, 25 $NaHCO_3$, 25 Dextrose, 3 Ascorbic Acid, 3.1 Sodium Pyruvate, 7 $MgCl_2$, 0.5 $CaCl_2.2H_2O$, and 0.2 Kyneuric Acid. The pH was kept at 7.3-7.4 whereas the osmolality was set to 280-285 mOsm. Slices were first prepared in the semi-frozen buffer, always bubbled with 95% $O_2$ /5% $CO_2$ and then transferred to nominal ACSF that contained (mM): 125 NaCl, 3 KCl, 1.25 $NaH_2PO_4$, 25 $NaHCO_3$, 10 Dextrose, 2 $MgCl_2$, and 2 $CaCl_2.2H_2O$ with 7.3-7.4 pH and 300-305 mOsm osmolality. In a few experiments, reduced $CaCl_2$ concentration (0.5 mM) was used to avoid precipitation of divalent ions and attenuate $Ca^{2+}$-dependent $K^+$ currents. ACSF was also saturated with 95% $O_2$/5% $CO_2$. The pipette (internal) solution consisted of (mM): 130 K-Gluconate, 6 KCl, 3 NaCl, 10 HEPES, 0.2 EGTA, 4 ATP-

Mg, 0.4 GTP-Tris, and 14 phosphocreatine-Tris. The pH was adjusted with KOH to 7.25 and osmolality to 300 mOsm. The recording ACSF was modified for different experiments as described in Supplementary Table S2.

For experiments depicted in figures 2 & 4 and Supplementary figures S5& S10, depolarizing steps in the multi-step protocol showed evidence of transiently activated depolarization-dependent $K^+$ currents since 4-AP or TEA-Cl were not included in the ACSF (Supplementary Table S2). These transient currents were not observed in all other multi-step experiments since compounds blocking depolarization-activated $K^+$ currents were continuously present (Supplementary Table S2).

*Experiments involving blockade of KCNK2 with acidification and Spadin*

For experiments in figure 3 (*Right*) targeting $K_{Leak}$, slices were kept in nominal ACSF. Only after patching the neurons, the actual control ACSF was applied for at least 3 min before data collection. In the same experiments, control ACSF contained arachidonic acid (AA) with pH 8.0 for the purpose of enhancing KCNK2 currents. However, in separate experiments, we did not find any detectable effects of AA and pH 8.0 when applied individually. Moreover, in previous studies, it is unclear how effectively acidification or Spadin alone blocks KCNK2-mediated current in neurons, and in separate experiments we also observed a weak blockade when either was used in isolation. Therefore, we used both acidification and Spadin together assuming this would enhance KCNK2 blockade. To maintain the desired pH, HEPES buffered ACSF was used for this experiment.

*Statistics*

For all data, sample number is the number of neurons, and data are stated in this order: WT control followed by *Foxp1* deletion groups. All data are plotted as mean ± standard error. Unless stated otherwise, 2-way ANOVA for repeated measures is performed with Geisser-Greenhouse correction followed by the Holm-Sidak test for the correction for multiple comparisons. When ANOVA analysis is not employed, we use a Mann-Whitney (MW) t-test.

In some instances, we compare two genotypic difference plots which were obtained under two different reagent conditions. In this case, we analyze the current densities from control conditions and in presence of reagent for one membrane potential only and perform a 2-way ANOVA (genotype x reagent) with repeated measures for the reagent where a significant interaction term indicated that the current density at that potential was different. Because IV-plots contain 9 genotypic comparisons (one for each membrane potential point), the criterion p-value for this interaction is modified with a Bonferroni correction based on 9 comparisons.

*Single-cell RNA sequencing*

We used single-cell RNA sequencing data from our previous study [5]. In brief, using the 10X Genomics platform, the cell-type specific transcriptome of the control and *Foxp1* deleted D1 and D2 SPNs from P9 mice (N=4, for each genotype) was generated and sequenced. Both D1 and D2 control SPNs were collected from *Foxp1*$^{CTL}$ mice whereas *Foxp1* deleted D1 and D2 SPNs were collected from D1 and D2 *Foxp1*$^{cKO}$ mice respectively. While this dataset includes gene expression from all cell types in the striatum, we only included SPNs for analysis this study.

We used raw UMI count data from a total of 11,355 SPNs from the three genotypes (*Foxp1*$^{CTL}$: 2,961, D1 *Foxp1*$^{cKO}$: 4,839 and D2 *Foxp1*$^{cKO}$: 3,555) for clustering analysis using Seurat v3 [20-22] R analysis pipeline. The dataset was log-normalized, scaled using a factor of 10,000 and regressed (*linear*) for covariates (number of UMIs per cell, percent mitochondrial content per cell, batches and number of cells per library). Further, the top 2,000 highly variable genes were used to compute principal components (*PCs*). Using Jackstraw analysis, 43 statistically significant PCs were used to identify clusters of cells within the data with Louvain algorithm as described in Seurat analysis pipeline [21-23]. Cellular clusters were visualized using Uniform Manifold Approximation and Projection (UMAP) [23,24] in two dimensions.

Annotating the clusters based on our previously published dataset [5], D1 and D2 SPNs across all three genotypes were used to generate dot plots to show relative mRNA expression of KCNJ and KCNK channels. The color of the dot represents the scaled averaged log normalized expression across cells in a given cell type whereas its size represents the scaled percentage of cells expressing the gene among the same cell type. Dot plots were prepared using the *DotPlot* function of Seurat v3 [20-22].

For differentially expressed genes, either D1 or D2 SPNs were grouped within each genotype and were compared across genotypes such as D1 *Foxp1*$^{cKO}$ compared to *Foxp1*$^{CTL}$ for D1 SPNs and D2 *Foxp1*$^{cKO}$ compared to *Foxp1*$^{CTL}$ for D2 SPNs accounting for averaged expression differences. A Poisson likelihood ratio test from the Seurat v3 analysis pipeline was used to identify significant gene expression changes (adj. p-value $<=0.05$, $\log_2 FC \geq 0.3$) across genotypes within groups of cells (D1 or D2 SPNs) instead of identified cellular clusters.

**References**

1 Araujo, D. J. *et al.* Foxp1 in Forebrain Pyramidal Neurons Controls Gene Expression Required for Spatial Learning and Synaptic Plasticity. *J Neurosci* **37**, 10917-10931, doi:10.1523/JNEUROSCI.1005-17.2017 (2017).

2 Feng, X. *et al.* Foxp1 is an essential transcriptional regulator for the generation of quiescent naive T cells during thymocyte development. *Blood* **115**, 510-518, doi:10.1182/blood-2009-07-232694 (2010).
3 Gong, S. *et al.* Targeting Cre recombinase to specific neuron populations with bacterial artificial chromosome constructs. *J Neurosci* **27**, 9817-9823, doi:10.1523/JNEUROSCI.2707-07.2007 (2007).
4 Ade, K. K., Wan, Y., Chen, M., Gloss, B. & Calakos, N. An Improved BAC Transgenic Fluorescent Reporter Line for Sensitive and Specific Identification of Striatonigral Medium Spiny Neurons. *Front Syst Neurosci* **5**, 32, doi:10.3389/fnsys.2011.00032 (2011).
5 Anderson, A. G., Kulkarni, A., Harper, M. & Konopka, G. Single-Cell Analysis of Foxp1-Driven Mechanisms Essential for Striatal Development. *Cell Rep* **30**, 3051-3066 e3057, doi:10.1016/j.celrep.2020.02.030 (2020).
6 Rajkovich, K. E. *et al.* Experience-Dependent and Differential Regulation of Local and Long-Range Excitatory Neocortical Circuits by Postsynaptic Mef2c. *Neuron* **93**, 48-56, doi:10.1016/j.neuron.2016.11.022 (2017).
7 Adesnik, H., Li, G., During, M. J., Pleasure, S. J. & Nicoll, R. A. NMDA receptors inhibit synapse unsilencing during brain development. *Proc Natl Acad Sci U S A* **105**, 5597-5602, doi:10.1073/pnas.0800946105 (2008).
8 Hays, S. A., Huber, K. M. & Gibson, J. R. Altered Neocortical Rhythmic Activity States in Fmr1 KO Mice Are Due to Enhanced mGluR5 Signaling and Involve Changes in Excitatory Circuitry. *The Journal of Neuroscience* **31**, 14223-14234, doi:10.1523/jneurosci.3157-11.2011 (2011).
9 Agmon, A. & Connors, B. W. Thalamocortical responses of mouse somatosensory (barrel) cortex in vitro. *Neuroscience* **41**, 365-379 (1991).
10 Marty, A. & Neher, E. in *Single-channel recording* (eds B. Sakmann & E. Neher) Ch. 2, 31-52 (Plenum Press, 1995).
11 Zar, J. H. *Biostatistical Analysis*. (Prentice Hall, 1999).
12 Gibson, J. R., Bartley, A. F. & Huber, K. M. Role for the subthreshold currents ILeak and IH in the homeostatic control of excitability in neocortical somatostatin-positive inhibitory neurons. *J Neurophysiol* **96**, 420-432 (2006).
13 Ketchum, K. A., Joiner, W. J., Sellers, A. J., Kaczmarek, L. K. & Goldstein, S. A. A new family of outwardly rectifying potassium channel proteins with two pore domains in tandem. *Nature* **376**, 690-695, doi:10.1038/376690a0 (1995).
14 Mazella, J. *et al.* Spadin, a sortilin-derived peptide, targeting rodent TREK-1 channels: a new concept in the antidepressant drug design. *PLoS Biol* **8**, e1000355, doi:10.1371/journal.pbio.1000355 (2010).
15 Shen, W., Hamilton, S. E., Nathanson, N. M. & Surmeier, D. J. Cholinergic suppression of KCNQ channel currents enhances excitability of striatal medium spiny neurons. *J Neurosci* **25**, 7449-7458, doi:10.1523/JNEUROSCI.1381-05.2005 (2005).
16 Coetzee, W. A. *et al.* Molecular diversity of K+ channels. *Ann N Y Acad Sci* **868**, 233-285 (1999).
17 Koyama, S. & Appel, S. B. Characterization of M-current in ventral tegmental area dopamine neurons. *J Neurophysiol* **96**, 535-543, doi:10.1152/jn.00574.2005 (2006).
18 Schindelin, J. *et al.* Fiji: an open-source platform for biological-image analysis. *Nat Methods* **9**, 676-682, doi:10.1038/nmeth.2019 (2012).
19 Schneider, C. A., Rasband, W. S. & Eliceiri, K. W. NIH Image to ImageJ: 25 years of image analysis. *Nat Methods* **9**, 671-675 (2012).
20 Butler, A., Hoffman, P., Smibert, P., Papalexi, E. & Satija, R. Integrating single-cell transcriptomic data across different conditions, technologies, and species. *Nat Biotechnol* **36**, 411-420, doi:10.1038/nbt.4096 (2018).

21 Stuart, T. *et al.* Comprehensive Integration of Single-Cell Data. *Cell* **177**, 1888-1902 e1821, doi:10.1016/j.cell.2019.05.031 (2019).

22 Satija, R. *Seurat - Guided Clustering Tutorial*, <https://satijalab.org/seurat/v3.1/pbmc3k_tutorial.html> (2019).

23 Becht, E. *et al.* Dimensionality reduction for visualizing single-cell data using UMAP. *Nature Biotechnology* **37**, 38-+, doi:10.1038/nbt.4314 (2019).

24 Leland McInnes, J. H., James Melville. *UMAP: Uniform Manifold Approximation and Projection for Dimension Reduction*, <https://arxiv.org/abs/1802.03426> (2018).
