## Supplementary Tables for "FOXP1 negatively regulates intrinsic excitability in D2 striatal projection neurons by promoting inwardly rectifying and leak potassium currents"

| Compound | Figure | Con (mM) | Target Chan | Action | Vendor | Ref |
|---|---|---|---|---|---|---|
| **TTX** | 3-6, S1,S3,S4 | 0.001 | Volt-Dep $Na^+$ | Block | hellobio | (Hille, 2001b) |
| **$CdCl_2$** | 3-6, S1,S3,S4 | 0.2 | Volt-Dep $Ca^{2+}$ , $K_{CA}$ | Block | Sigma | (Hille, 2001c) |
| **4-AP** | 4-5 | 2.5 | $K_{DR}$, $K_A$, $K_D$ | Block | Sigma | (Amarillo et al., 2008) |
| **TEA-Cl** | 4-5 | 5.0 | $K_{DR}$, $K_A$, $K_{CA}$, KCNQ (50-80%) | Block | Sigma | (Amarillo et al., 2008), (Hibino et al., 2010; Shen et al., 2005) |
| **CsCl** | 3-6, S3 | 1.0 | $K_{IR}$, $K_{DR}$, $K_A$, $K_{CA}$, $I_H$ | Block | Sigma | (Lieberman et al., 2018; Mermelstein et al., 1998) |
| **$BaCl_2$** | 4,5 | 4.0 | $K_{IR}$, KCNK2, other $K_{Leak}$, $K_{DR}$, KCNQ | Block | Sigma | (Lieberman et al., 2020; Ma et al., 2011; Shen et al., 2007) |
| **Spadin** | 5 | 0.0005 | KCNK2 ($K_{Leak}$) | Inhib | Tocris | (Djillani et al., 2017; Mazella et al., 2010) |
| **Low pH** | 5 | pH 6.2 | KCNK2 ($K_{Leak}$) | Inhib | | (Goldstein et al., 2001; Sandoz et al., 2009) |

**Supplementary Table S1.** Compounds used in experiments. Additional general references for all compounds (Coetzee et al., 1999; Hille, 2001a; Johnston and Wu, 1995).

| | Measurement | Figure | Control ACSF | Wash-in reagents | Control ACSF blockade |
|---|---|---|---|---|---|
| **1.** | Excitability | 1A,B,G,H,I<br>2A,B<br>6A,B,H,I<br>S2A,B<br>S5A,B | nACSF | | |
| **2.** | $Cs^+$-sensitive currents ($K_{IR}$) | 3,6E-G,S3 | nACSF with 0.2 $CdCl_2$ and 0.001 TTX | 1.0 CsCl to block $K_{IR}$ | $CdCl_2$ and TTX block $Ca^{2+}$ and $Na^+$ channels, respectively. |
| **3.** | $Ba^{2+}$-sensitive currents ($K_{Leak}$) | 4 | Same as (2) with added 5 TEA-Cl, 2.5 4-AP and 1.0 CsCl. CaCl2 reduced to 0.5 to avoid precipitation. | 4.0 $BaCl_2$ to block $K_{Leak}$. | Same as (2) with blockade of depolarization-dependent $K^+$ channels. |
| **4.** | $K_{Leak}$ and KCNK2 | 5 | HEPES ACSF with 0.001 TTX, 0.01 AA, 0.2 $CdCl_2$, 1.0 CsCl, 5.0 TEA-Cl, 2.5 4-AP, 4.0 $BaCl_2$, pH 8.0 | pH 6.20 and 500 nM Spadin to block KCNK2 | Same as (3) with additional partial blockade of $K_{IR}$ and $K_{Leak}$ and blockade of KCNQ. |

**Supplementary Table S2.** Modifications to ACSF for different experiments. AA=Arachidonic Acid. Compound concentrations in mM.

| | -55 mV membrane potential | | Normalized conductance, G (pS/pF) | |
|---|---|---|---|---|
| | | **Figure** | ***Foxp1*** $^{CTL}$ | ***Foxp1*** $^{cKO}$ |
| **1** | Control | 2C, *left* | 107 | 62 |
| **2** | Cs + 4-AP + TEA | 3A, *left* | 84 | 51 |
| **3** | Cs + 4-AP + TEA + Ba | 3A, *right* | 29 | 24 |
| **4** | [2] – [3] = $K_{Leak}$ | | 55 | 27 |
| **5** | [4]/[1] = % of total G from $K_{Leak}$ | | 51% | 42% |
| **6** | Down-regulated $K_{Leak}$ G with *Foxp1* deletion based on pharmacology, CTL – cKO in [4] | | | 28 |
| **7** | Down-regulated G with *Foxp1* deletion, [1] | | | 45 |
| **8** | % down-regulated G accounted for by pharm, $K_{Leak}$, [6]/[7] | | | 62% |

**Supplementary Table S3. Calculations of the $K_{Leak}$ contribution to the normalized conductance.** Based on normalized conductance data obtained at -55 mV membrane potential (see Suppl. Methods, *Membrane potentials for measuring $K_{IR}$ and $K_{Leak}$*), we calculated the approximate contribution of a putative $K_{Leak}$ to the total normalized conductance of *Foxp1*$^{CTL}$ D2 SPNs as well as the amount that is regulated by *Foxp1*. We used -55 mV data from $Cs^+$ (Fig. 2C) and $Ba^{2+}$ wash-in experiments (Fig. 3A) to make this calculation. Assuming that the $Ba^{2+}$-sensitive conductance reflects $K_{Leak}$ (Fig. 3A), we find that the $K_{Leak}$ accounted for 51% of the total conductance in normally developing D2 SPNs. Even with the possibility that our calculations involve some contamination by other currents, these numbers suggest a significant impact on excitability by $K_{Leak}$. Deletion of *Foxp1* in D2 SPNs reduced the contribution of $K_{Leak}$ to 42% of the total conductance (Fig. 3A). There was a decrease of 45 pS/pF in the total conductance of D2 SPNs with *Foxp1* deletion, where almost 62% (28 pS/pF) was due to a downregulation in $K_{Leak}$.
